# Making the MOSTest of imaging genetics

**DOI:** 10.1101/767905

**Authors:** Dennis van der Meer, Oleksandr Frei, Tobias Kaufmann, Alexey A. Shadrin, Anna Devor, Olav B. Smeland, Wes Thompson, Chun Chieh Fan, Dominic Holland, Lars T. Westlye, Ole A. Andreassen, Anders M. Dale

## Abstract

Regional brain morphology has a complex genetic architecture, consisting of many common polymorphisms with small individual effects, which has proven challenging for genome-wide association studies to date, despite its high heritability^1,2^. Given the distributed nature of the genetic signal across brain regions, joint analysis of regional morphology measures in a multivariate statistical framework provides a way to enhance discovery of genetic variants with current sample sizes. While several multivariate approaches to GWAS have been put forward over the past years^3–5^, none are optimally suited for complex, large-scale data. Here, we applied the Multivariate Omnibus Statistical Test (MOSTest), with an efficient computational design enabling rapid and reliable permutation-based inference, to 171 subcortical and cortical brain morphology measures from 26,502 participants of the UK Biobank (mean age 55.5 years, 52.0% female). At the conventional genome-wide significance threshold of α=5×10^−8^, MOSTest identifies 347 genetic loci associated with regional brain morphology, more than any previous study, improving upon the discovery of established GWAS approaches more than threefold. Our findings implicate more than 5% of all protein-coding genes and provide evidence for gene sets involved in neuron development and differentiation. As such, MOSTest, which we have made publicly available, enhances our understanding of the genetic determinants of regional brain morphology.

---

Regional surface area and thickness of the cerebral cortex and volume of subcortical structures are highly heritable brain morphological features with complex genetic architectures, involving many common genetic variants with small effect sizes^1,2^. The predominant strategy for identifying genomic loci associated with complex traits is through genome-wide association studies (GWAS), a mass-univariate approach whereby the association between a single outcome measure and each of millions of genetic variants, in isolation, is tested. This is accompanied by a stringent multiple comparison correction to control the family-wise error rate, necessitating very large sample sizes to identify even relatively strong effects. To date, the largest GWAS of regional brain morphological features, based on brain scans obtained from up to fifty thousand individuals, identified almost two hundred genetic variants^1^, which together explained only a fraction of the reported narrow-sense heritability. These studies primarily investigate each region of interest individually, compounding the multiple comparisons correction problem.

In addition to small effect sizes across many variants, the genetic architectures of sets of regional brain features are likely to strongly overlap. Gene expression studies of the human brain have shown widespread gradients across the cortex^6^. Thus, genetic variants probably have distributed effects across regions and morphological measures. While cortical thickness and surface area have been reported to be phenotypically and genetically only weakly correlated to each other^7^, many brain-related traits, e.g. mental disorders, share a large proportion of genetic variants, even in the absence of an overall correlation^8^. The discovery of these variants may be boosted through joint analysis of these traits, in a multivariate framework. This avoids the family-wise error rate penalty for studying multiple outcome measures, or the use of strategies that reduce phenotypic information to a single composite score, which can cause considerable loss of statistical power ^9^. Importantly, a multivariate approach is much more consistent with the notion of the brain being an integrated unit, with highly interconnected and biologically similar brain regions, compared to univariate approaches that ignore the information shared across these component measures.

Here we implement a Multivariate Omnibus Statistical Test (MOSTest), designed to boost the power of imaging genetics by capitalizing on the distributed nature of genetic influences across brain regions and pleiotropy across imaging modalities. MOSTest is efficient and capable of combining large-scale genome-wide analyses of dozens of measures for tens of thousands of individuals within hours while achieving enhanced statistical power. Key steps of the MOSTest analysis include: 1) applying a rank-based inverse normal transformation to the input measures; 2) estimating the multivariate correlation structure from the GWAS on randomly permuted genotype data; 3) calculating the Mahalanobis norm, as the sum of squared de-correlated z-values across univariate GWAS summary statistics, to integrate effects across the measures into a multivariate test statistic; and 4) employing the gamma cumulative density function to fit an analytic form for the null distribution, enabling extrapolation to and beyond the 5×10^−8^ significance threshold. This avoids the extensive computational burden associated with permutation-based approach which has, until now, prohibited its application in GWAS. The Supplementary Materials contains a detailed description and validation of these steps through simulations.

We compare the MOSTest with an established inferential methodology recently used by the Enhancing NeuroImaging Genetics through Meta-Analysis (ENIGMA) consortium^1^, referred to as the min-P approach; min-P takes the smallest p-value of each SNP across the univariate GWAS, and corrects this for the effective number of traits studied^4,10^, i.e. shared genetic architecture across traits does not contribute to statistical power. Min-P achieves its maximum power when the genetic effects across traits are independent; conversely, multivariate approaches have greater power when genetic effects are distributed across traits ^3^.

Here, we applied MOSTest to sets of regional brain morphology measures, hypothesizing that it will outperform the min-P approach due to the distributed nature of genetic effects and the presence of pleiotropy across modalities. Our sample consisted of 26,502 healthy White European participants of the UK Biobank (UKB), with a mean age of 55.5 years (standard deviation (SD) 7.4 years), 52.0% female. We processed T_1_-weighted structural MRI scans with the FreeSurfer v5.3 standard “recon-all” processing pipeline, producing a subset of 35 subcortical volume estimates ^11^, as well as subsets of surface area and cortical thickness estimates, both consisting of 68 cortical regions following the Desikan-Killiany parcellation^12^, for a total set of 171 measures. All measures were pre-residualized for age, sex, scanner site, a proxy of image quality (FreeSurfer’s Euler number), the first twenty genetic principal components to control for population stratification, and a global measure specific to each set of variables: mean cortical thickness for the regional thickness measures, total surface area for the area measures, and intracranial volume for the subcortical structures. Subsequently, we performed a rank-based inverse normal transformation to the residualized measures. We made use of the UKB v3 imputed data, carrying out standard quality-checks and setting a minor allele frequency threshold of .005, leaving 7.4 million SNPs. Univariate GWAS of each measure was performed with standard tools. The resulting summary statistics were then tested for overall significance with MOSTest. Independent significant SNPs and genomic loci were identified in accordance with FUMA SNP2GENE definition^13^. Details on sample composition, processing of the data, and the analysis techniques are presented in the Online Methods section.

The multivariate GWAS identified thousands of independent SNPs reaching the genome-wide significance threshold of α=5×10^−8^ across hundreds of independent loci, as shown in Figure 1A. See the Extended Data section for the list of discovered loci per feature set. Overall, MOSTest led to a threefold higher discovery than the min-P approach. The difference in performance is particularly pronounced when all features are combined, as is also evident from the Miami plots shown in Figure 1B through E. As can be seen in Figure 1F, many loci identified with MOSTest were shared across the three feature subsets. Further, the number of MOSTest-discovered loci for all features combined (347) is larger than the summation of number of unique loci found when analyzing the feature subsets individually (301). This illustrates how MOSTest capitalizes on the distributed and non-sparse nature of genetic effects through the combination of features.

**Figure 1.**
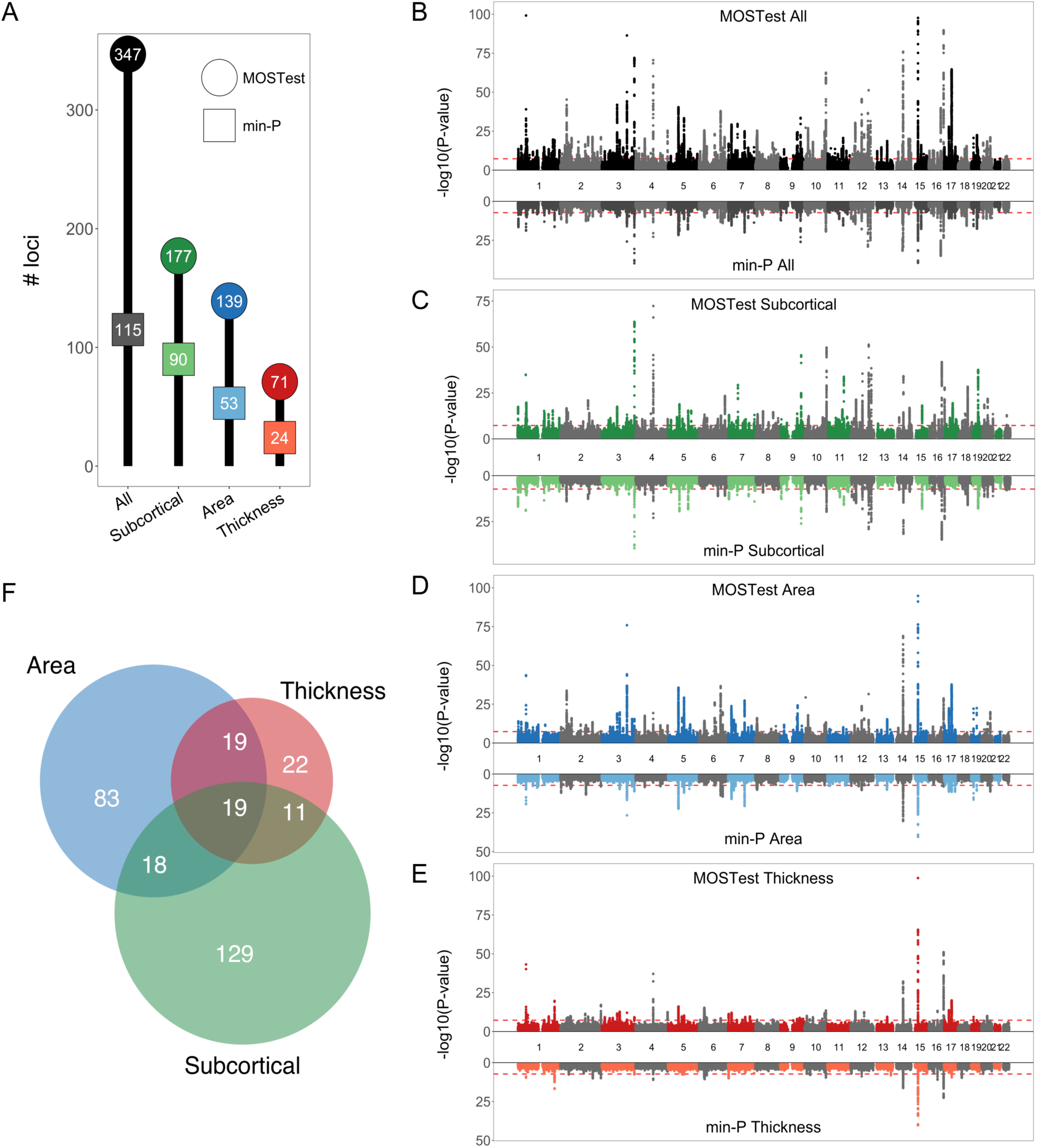
Highly improved locus discovery through MOSTest. **A.** Number of independent whole-genome significant loci identified (on the y-axis and in the bubbles) for each set of features (on the x-axis), by MOSTest (in darker colored circles) and by min-P (in lighter colored squares). **B - E.** Miami plots, contrasting the observed -log_10_(p-values), shown on the y-axis, of each SNP for MOSTest (top half) with min-P (bottom half), for each of the feature sets. The x-axis shows the relative genomic location, grouped by chromosome, and the red dashed lines indicate the whole-genome significance threshold of 5×10^−8^. Note, y-axis is clipped at -log_10_(p-value)=100. **F.** Venn diagram depicting the number of loci, identified through MOSTest, overlapping between the three feature subsets.

We carried out replication analyses of the GWAS results in an additional 4,884 UKB participants, whose neuroimaging data was released after we carried out our primary analyses. The loci discovered through MOSTest and min-P replicated at similar levels, with approximately 40% being significant in this smaller additional sample. Therefore, the absolute number of loci replicating is three times higher for MOSTest compared to min-P. The full results are shown in the Extended Data.

We performed extensive validation of MOSTest methodology and implementation, confirming that it has correct type-I error in simulations with synthetic data (Figure S4) and on real data under permutations (Figure S2). We also carried out a formal comparison between the MOSTest and MV-PLINK^5^, finding that the two methods have similar statistical power for detecting associations (Figure S1), while the MOSTest analysis ran 10,000 times faster. Further, we performed LD score regression and used the intercept to validate that MOSTest results are free of genomic inflation (Table S5).

Using the MiXeR tool ^8,14^ we fitted a Gaussian mixture model of the null and non-null effects to the GWAS summary statistics, estimating for each feature set the number of SNPs involved, i.e. their combined polygenicity, and their effect size variance, or ‘discoverability’. Please see the Online Methods for more details. The results are summarized in Figure 2, depicting the estimated proportion of genetic variance explained by discovered SNPs by both approaches as a function of sample size. The horizontal shift of the curve indicates that the effective sample size of MOSTest is generally about twice as high as that of min-P, with the highest discovery for MOSTest when all features are combined, and lowest discovery for the set of cortical thickness features.

**Figure 2.**
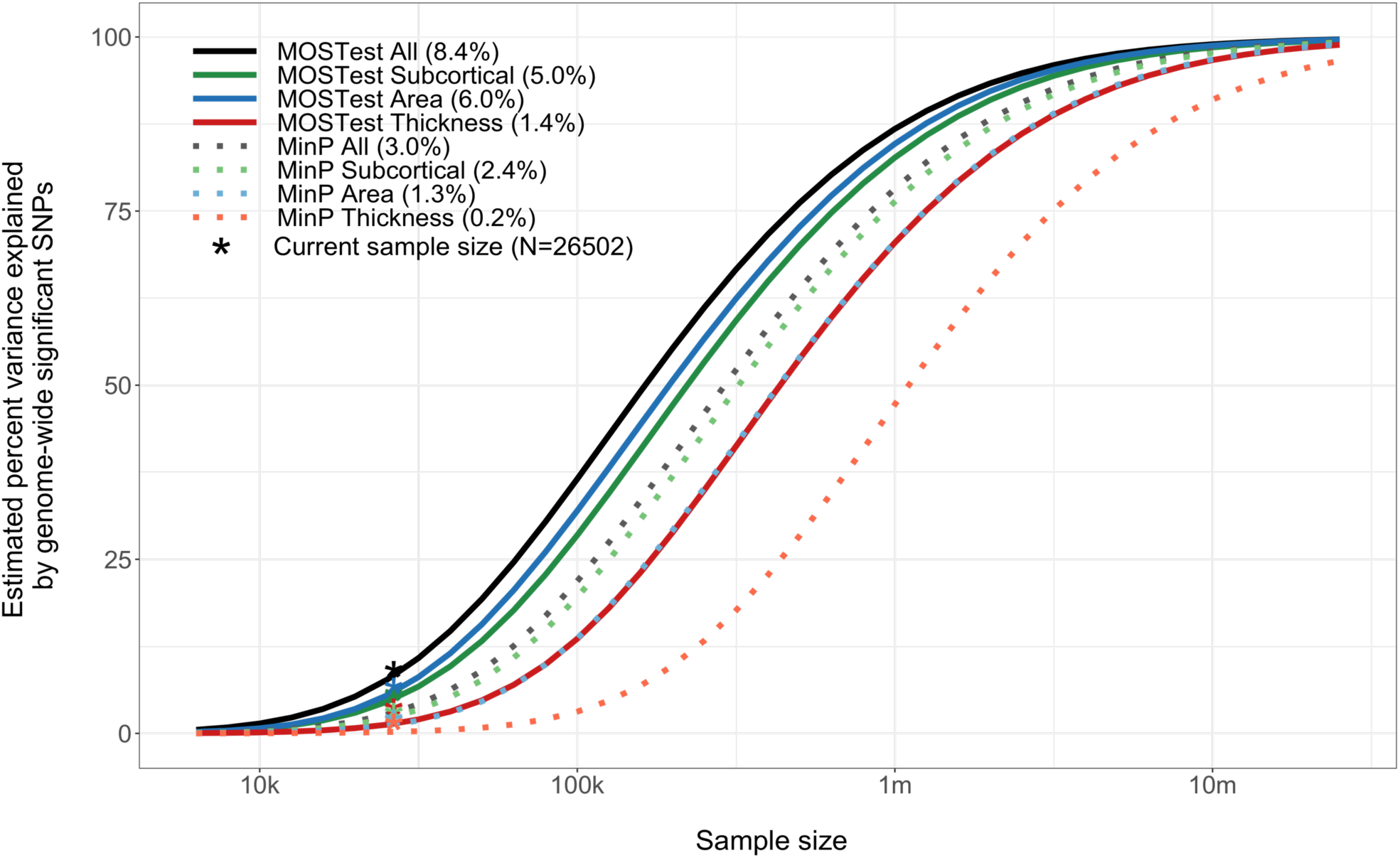
MOSTest increases effective sample size. Estimated percent of genetic variance explained by SNPs surpassing the genome-wide significance threshold, on the y-axis, as a function of sample size, depicted on the x-axis on a log_10_ scale, for each of the feature sets and for both approaches. Percentages of genetic variance explained by discovered SNPs with current sample size (N=26,502) are shown in parentheses.

Cortical maps, depicting the morphological associations of the lead SNPs identified with MOSTest on all features with regional surface area and thickness measures, made clear that these SNPs have distributed effects, often with mixed directions, across regions and feature sets. As an example, Figure 3 shows the maps for the top two hits (rs1080066 on chromosome 15, p=1.2×10^−305^, and rs13107325 on chromosome 4, p=3.1×10^−124^), all other maps are available in the Supplementary Material. These maps revealed anterior-posterior gradients as well as hemisphere-specific effects of some of the lead SNPs, in line with reported genetic patterns of the brain ^15,16^.

**Figure 3.**
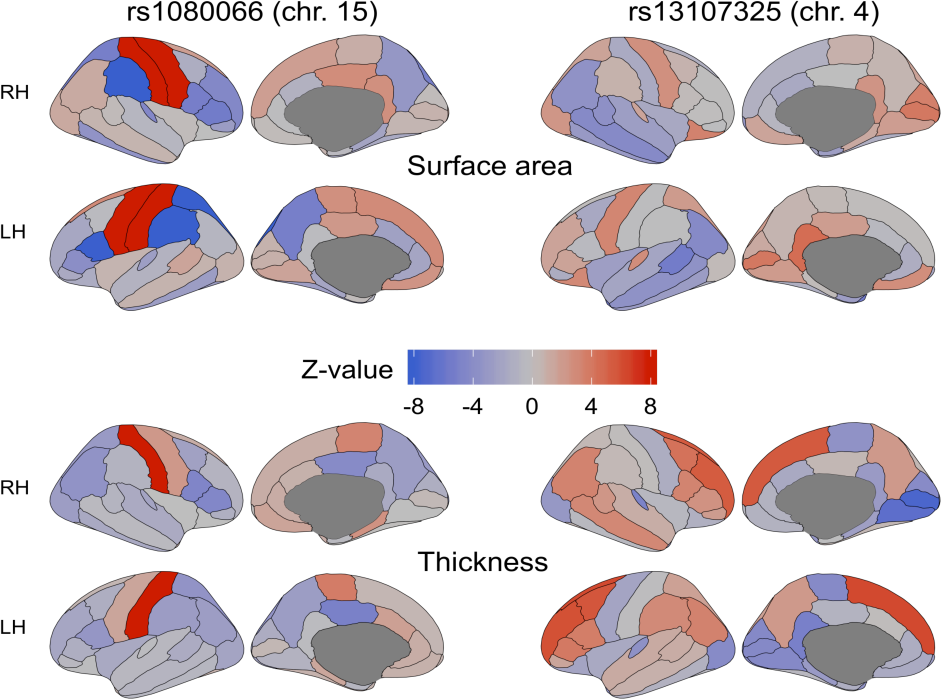
The genetic variants identified through MOSTest have distributed effects across the cortex. Z-values from the univariate GWAS for each cortical region for the two most significant lead SNPs from MOSTest applied to all features combined (left two columns for rs1080066 on chromosome 15, and right two columns for rs13107325 on chromosome 4). The top two rows show the effects of the SNPs on regional surface area, and the bottom two on cortical thickness. Positive effects of carrying the minor allele are shown in red, and negative in blue. Note: the absolute Z-value scaling is clipped at 8 (p=1.2*10^−15^), RH=right hemisphere, LH=left hemisphere.

Gene-level analyses, using Multi-marker Analysis of GenoMic Annotation (MAGMA) 13,17, indicated that 1034 out of all 18,775 protein coding genes (i.e. 5.5%) were significant, with a p-value below a Bonferroni corrected threshold of □=.05/18,775. Figure 4A shows the number of significant genes for each set of features. Through competitive gene-set analyses we identified 136 significant Gene Ontology sets for MOSTest applied to all features, the vast majority of which related to neuronal development and differentiation, with Figure 4B listing the top fifteen.

**Figure 4.**
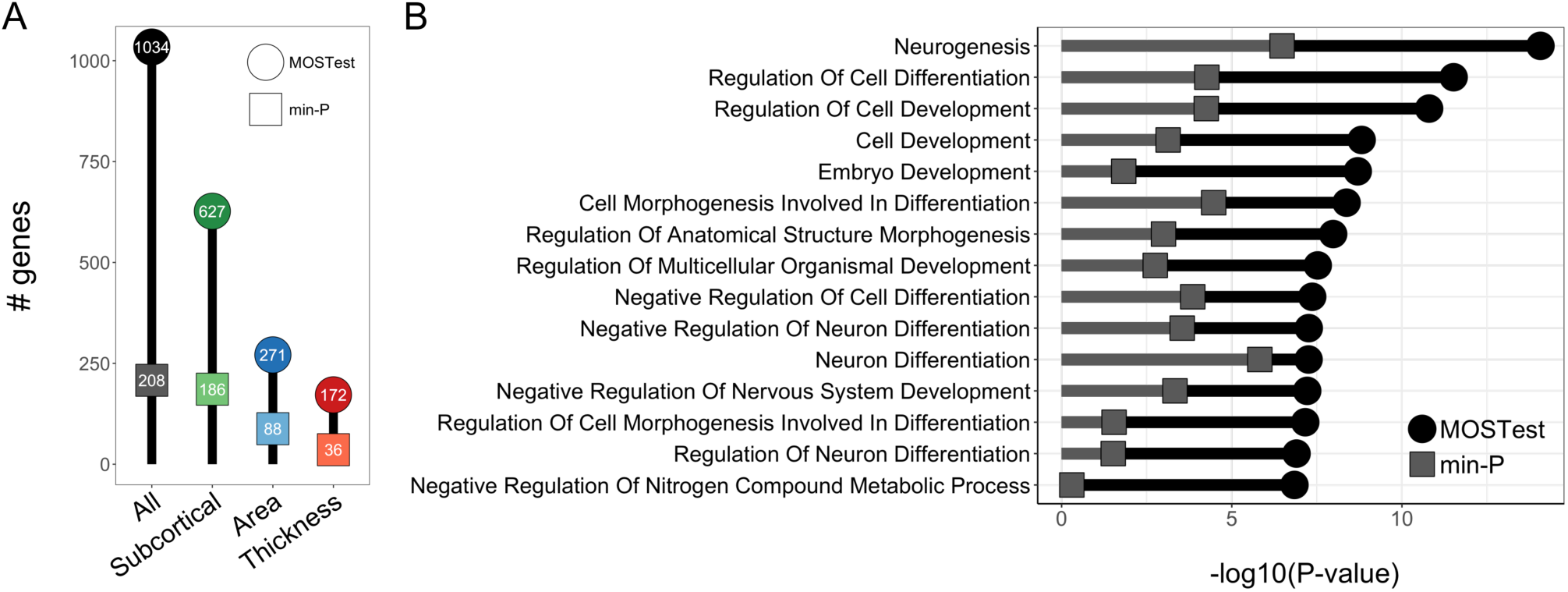
Functional mapping and annotation indicates high neurobiological relevance of our findings. **A.** Number of whole-genome significant genes identified (on the y-axis and in the bubbles) for each set of features (on the x-axis), by MOSTest (in darker colored circles) and by min-P (in lighter colored squares). **B.** Results from the gene-set analyses following the application of the multivariate GWAS on all brain features. The fifteen most significant Gene Ontology sets for MOSTest are listed on the y-axis and -log10(p-values) are shown on the x-axis. MOSTest Bonferroni corrected p-values are indicated by black circles, min-P in grey squares.

Applying the MOSTest approach to structural brain imaging data, we discovered more loci associated with regional cortical and subcortical morphology than any previous GWAS of brain morphology, even those that had nearly double the sample size^1,2^. Further, a direct comparison with established methods in the same sample revealed a threefold increase in discovery. This improvement indicates the presence of extensive shared genetic architecture across brain regions and across morphological measures, attesting to the importance of estimating levels of polygenic overlap beyond those indicated by genetic correlations^8^, and arguing for techniques that boost discovery of genetic determinants leveraging shared signal between traits^18^. Indeed, overlapping genetic determinants are to be expected given the shared genetic control of neurodevelopment across brain regions^19^, and that similar molecular mechanisms operate across regional borders defined by gross morphological features. This is in accordance with the high levels of pleiotropy across many brain-related traits and disorders^20^. Therefore, our multivariate strategy is better tailored to the underlying neurobiological processes than conventional univariate approaches, as confirmed by our identification of highly significant links to gene sets of neuronal development and differentiation. Further, our extensive validation of the MOSTest methodology and implementation show that the MOSTest is a valid statistical test with an improved statistical power to detect associations in a multivariate context, suitable for applications to large-scale data.

With the large gain of power and consequently lower required sample sizes of MOSTest, we predict that it will be possible to uncover the majority of SNPs influencing brain morphology in the upcoming years. The UKB initiative, for instance, is set to release neuroimaging data of a 100,000 individuals by 2022^21^, which we estimate will enable MOSTest to identify SNPs that will explain about 40% of the additive genetic variance in regional brain morphology. The output of MOSTest is suited for secondary analyses and follow-up studies to investigate the relation between the set of loci discovered and individual features, with a much decreased multiple-comparisons burden. The MOSTest code is publicly available, see the Supplementary Materials. In addition to brain structure, the MOSTest approach may also be of value in uncovering the genetic determinants of brain function and other complex human phenotypes consisting of correlated component measures, such as mental, cognitive or cardiometabolic phenotypes, by taking advantage of the rich multivariate datasets now available.

## Supporting information

Brain maps MOSTest All

## Author contributions

A.M.D., O.F., D.v.d.M and O.A.A. conceived the study; D.v.d.M., O.F., T.K. and A.M.D. pre-processed the data. D.v.d.M., O.F. and A.M.D. performed all analyses, with conceptual input from O.A.A.; All authors contributed to interpretation of results; D.v.d.M. drafted the manuscript and all authors contributed to and approved the final manuscript.

## Materials & Correspondence

The data incorporated in this work were gathered from public resources. The code will be made available via https://github.com/precimed/mostest (GPLv3 license) upon publication of this study. Correspondence and requests for materials should be addressed to

## Acknowledgements

The authors were funded by the Research Council of Norway (276082, 213837, 223273, 204966/F20, 229129, 249795/F20, 225989, 248778, 249795), the South-Eastern Norway Regional Health Authority (2013-123, 2014-097, 2015-073, 2016-064, 2017-004), Stiftelsen Kristian Gerhard Jebsen (SKGJ-Med-008), The European Research Council (ERC) under the European Union’s Horizon 2020 research and innovation programme (ERC Starting Grant, Grant agreement No. 802998) and National Institutes of Health (R01MH100351, R01GM104400). This work was partly performed on the TSD (Tjeneste for Sensitive Data) facilities, owned by the University of Oslo, operated and developed by the TSD service group at the University of Oslo, IT-Department (USIT). Computations were also performed on resources provided by UNINETT Sigma2 - the National Infrastructure for High Performance Computing and Data Storage in Norway.

## Competing financial interests

Dr. Andreassen has received speaker’s honorarium from Lundbeck, and is a consultant to HealthLytix. Dr. Dale is a Founder of and holds equity in CorTechs Labs, Inc, and serves on its Scientific Advisory Board. He is a member of the Scientific Advisory Board of Human Longevity, Inc. and receives funding through research agreements with General Electric Healthcare and Medtronic, Inc. The terms of these arrangements have been reviewed and approved by UCSD in accordance with its conflict of interest policies. The other authors declare no competing financial interests.

## Online Methods

### Sample

We made use of data from participants of the UKB population cohort, obtained from the data repository under accession number 27412. The composition, set-up, and data gathering protocols of the UKB have been extensively described elsewhere^22^. For this study, we selected White Europeans that had undergone the neuroimaging protocol. For the primary analysis, making use of T1 MRI scan data released up to April 2019, we excluded 1094 individuals with a primary or secondary ICD10 diagnosis of a neurological or mental disorder, as well as 594 individuals with bad structural scan quality as indicated by an age and sex-adjusted Euler number^23^ more than three standard deviations lower than the scanner site mean. Our sample size for this analysis was n=26502, with a mean age of 55.51 years (SD=7.42). 51.97% of the sample was female. For the replication analyses, we made use of an additional neuroimaging batch released in September 2019. After identical preprocessing steps as the primary sample, this sample consisted of 4,884 individuals with a mean age of 55.47 years (SD=7.37), 52.42% was female.

### Data preprocessing

T_1_-weighted scans were collected from three scanning sites throughout the United Kingdom, all on identically configured Siemens Skyra 3T scanners, with 32-channel receive head coils. The UKB core neuroimaging team has published extensive information on the applied scanning protocols and procedures, which we refer to for more details^21^. The T_1_ scans were obtained from the UKB data repositories and stored locally at the secure computing cluster of the University of Oslo. We applied the standard “recon-all -all” processing pipeline of Freesurfer v5.3, performing automated surface-based morphometry and subcortical segmentation^11,12^. From the output, we extracted the sets of regional subcortical and cortical morphology measures, as well as estimated intracranial volume (eICV). Table S1 contains all the regional morphology measures, per subset, included in the current study. For each of these, we included both the left and right hemisphere measure, if applicable.

We subsequently regressed out age, sex, scanner site, Euler number, and the first twenty genetic principal components from each measure. We further regressed out a global measure specific to each of the feature subsets: eICV for the subcortical volumes, mean thickness for the regional thickness measures, and total surface area for the regional surface area measures. This was done to ensure we are studying the genetic determinants of regional brain morphology rather than global effects. Following this, we applied rank-based inverse normal transformation^24^ to the residuals of each measure: : 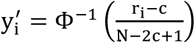, where r is the ordinary rank of the measure for i-th individual, N gives the sample size, Φ^−1;^ denotes the standard normal quantile, 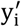 is the value after transformation, and c=0.5. This leads to normally distributed input into the univariate GWAS. See the Extended Data section below for a more in-depth discussion of the importance of this normalization procedure.

### Univariate GWAS procedure

We made use of the UKB v3 imputed data, which has undergone extensive quality control procedures as described by the UKB genetics team^25^. After converting the BGEN format to PLINK binary format, we additionally carried out standard quality check procedures, including filtering out individuals with more than 10% missingness, SNPs with more than 5% missingness, and SNPs failing the Hardy-Weinberg equilibrium test at p=1*10^−9^. We further set a minor allele frequency threshold of 0.005, leaving 7,428,630 SNPs.

The univariate GWAS on each of the 171 pre-residualized and normalized regional brain morphology measures were carried out using the standard additive model of linear association between genotype vector, g_j_, and phenotype vector, y. To speed up calculations we implemented the association test directly in MOSTest software. Statistical significance was assessed from Pearson’s correlation coefficient r_j_ = corr(y, g_.j_), as implemented in MATLAB’s corr function. This is equivalent to testing significance of the regression slope, 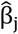, as both 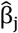 and r_j_ are assumed to be t-distributed and have the same t-value: 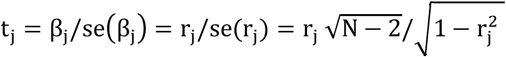, and therefore the same p-value, equal to Student’s t cumulative distribution function (cdf) with N − 2 degrees of freedom: P_val,j_ = 2 tcdf(− |t_j_|, N − 2), where N is the sample size^14^. Further, we validated that the above procedure produces the same results as the association test implemented in the commonly used PLINK’s additive model, option plink --assoc.

Independent significant SNPs and genomic loci were identified in accordance with FUMA SNP2GENE definition^13^. First, we select a subset of SNPs that pass genome-wide significance threshold 5×10^−8^ (calculated by minP or MOSTest), and use PLINK to perform a clumping procedure at LD r2=0.6, to identify the list of independent significant SNPs. Second, we clump the list of independent significant SNPs at LD r2=0.1 threshold to identify lead SNPs. Third, we query the reference panel for all candidate SNPs in LD r2 of 0.1 or higher with any lead SNPs. Further, for each lead SNP, it’s corresponding genomic loci is defined as a contiguous region of the lead SNPs’s chromosome, containing all candidate SNPs in r2=0.1 or higher LD with the lead SNP. Finally, adjacent genomic loci are merged together if they are separated by less than 250 KB. Allele LD correlations are computed from EUR population of the 1000 genomes Phase 3 data.

### MOSTest procedure

Let z_ij_ be the value of signed test statistic (z-score) calculated from the univariate association test between j-th SNP and i-th phenotype. Let z_j_ = (z_1j_, …, z_Kj_) be the vector of z-scores of j-th SNP across K phenotypes. Let Z = {z_ij_} be the matrix of z-scores, with rows corresponding to SNPs, and columns corresponding to phenotypes. Further, let 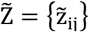 be the matrix of z-scores, calculated from association tests on a randomly permuted genotype vector of each SNP. To preserve correlation structure among phenotypes, the permutation was performed only once for each SNP, and the resulting genotype vector was used in association test across all phenotypes.

The MOSTest test statistic, 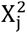, for the j-th SNP is calculated as Mahalanobis norm 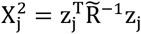, where 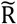 is the KxK correlation matrix of 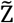. The null hypothesis of the MOSTest is that z_j_ is distributed as a multivariate normal random variable with zero mean and covariance 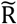. To compute the theoretical (i.e., under null) p-value of the MOSTest test statistic, we calculated the tail probability that a Chi-square statistics exceeds 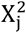. This probability is given by chi-square distribution with N degrees of freedom, or, equivalently, a gamma distribution, Gamma(K/2,0.5)^26^. Instead of using theoretical values, we fit the two free parameters of the Gamma(a,b) distribution to the observed distribution of 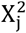 under permutation (shown in Table S4). The p-value of the MOSTest test statistic is then obtained from a cumulative distribution function of the gamma distribution, 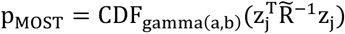.

Controlling for covariates, such as genetic principal components, is done via pre-residualization of all phenotype vectors, i.e. we replace them with the corresponding residual after multiple linear regression of the phenotype vector on the covariates. Additionally, we perform a rank-based inverse normal transformation of the residualized phenotypes, to ensure that z-scores forming the input to MOSTest are normally distributed.

MOSTest code will be made publicly available through GitHub upon publication of this article, https://github.com/precimed/mostest.

### MiXeR analysis

We applied a causal mixture model^8,14^ to estimate the percentage of variance explained by genome-wide significant SNPs as a function of sample size. For each SNP, i, MiXeR models its additive genetic effect of allele substitution, β_i_, as a point-normal mixture, 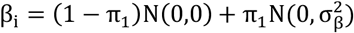, where π_1_ represents the proportion of non-null SNPs (‘polygenicity’) and 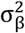 represents variance of effect sizes of non-null SNPs (‘discoverability’). Then, for each SNP, j, MiXeR incorporates LD information and allele frequencies for 9,997,231 SNPs extracted from 1000 Genomes Phase3 data to estimate the expected probability distribution of the signed test statistic, 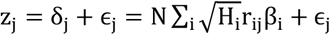, where N is sample size, H_i_ indicates heterozygosity of i-th SNP, r_ij_ indicates allelic correlation between i-th and j-th SNPs, and 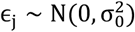 is the residual variance. Further, the three parameters, 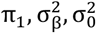, are fitted by direct maximization of the likelihood function. Fitting the univariate MiXeR model does not depend on the sign of z_j_, allowing us to calculate |z_j_| from MOSTest p-values. Finally, given the estimated parameters of the model, the power curve S(N) is then calculated from the posterior distribution p(δ_j_|z_j_, N).

### Gene-set analyses

We made use of the Functional Mapping and Annotation of GWAS (FUMA) online platform (https://fuma.ctglab.nl/) to further process the output from MOSTest and min-P. Through FUMA, we carried out MAGMA-based gene analyses using default settings, which entail the application of a SNP-wide mean model and use of the 1000 Genomes Phase 3 EUR reference panel. Gene-set analyses were done in a similar manner, restricting the sets under investigation to those that are part of the Gene Ontology biological processes subset (n=4436), as listed in the Molecular Signatures Database (MsigdB) v5.2.

## Extended Data

### Morphological features included

Please see Table S1 for an overview of all regional morphological features included in the analyses. We included all features outputted by the default Freesurfer subcortical and cortical processing streams, except for the range of global measures, CSF, surface holes, vessels, optic chiasm and hypointensities, as we did not consider these measures of regional brain morphology.

**Extended Data Table 1.** Regional brain morphology outcome measures, per subset, included in the study.

| *Subcortical Volumes* | *Surface Area* | *Thickness* |
| --- | --- | --- |
| Lateral Ventricle | Bankssts | Bankssts |
| Inferior Lateral Ventricle | Caudalanteriorcingulate | Caudalanteriorcingulate |
| Cerebellum White Matter | Caudalmiddlefrontal | Caudalmiddlefrontal |
| Cerebellum Cortex | Cuneus | Cuneus |
| Thalamus Proper | Entorhinal | Entorhinal |
| Caudate | Fusiform | Fusiform |
| Putamen | Inferiorparietal | Inferiorparietal |
| Pallidum | Inferiortemporal | Inferiortemporal |
| 3rd Ventricle* | Isthmuscingulate | Isthmuscingulate |
| 4th Ventricle* | Lateraloccipital | Lateraloccipital |
| Brain Stem | Lateralorbitofrontal | Lateralorbitofrontal |
| Hippocampus | Lingual | Lingual |
| Amygdala | Medialorbitofrontal | Medialorbitofrontal |
| Accumbens Area | Middletemporal | Middletemporal |
| Ventral Diencephalon | Parahippocampal | Parahippocampal |
| Choroid Plexus | Paracentral | Paracentral |
| 5th Ventricle* | Parsopercularis | Parsopercularis |
| CC Posterior* | Parsorbitalis | Parsorbitalis |
| CC Mid Posterior* | Parstriangularis | Parstriangularis |
| CC Central* | Pericalcarine | Pericalcarine |
| CC Mid Anterior* | Postcentral | Postcentral |
| CC Anterior* | Posteriorcingulate | Posteriorcingulate |
|  | Precentral | Precentral |
|  | Precuneus | Precuneus |
|  | Rostralanteriorcingulate | Rostralanteriorcingulate |
|  | Rostralmiddlefrontal | Rostralmiddlefrontal |
|  | Superiorfrontal | Superiorfrontal |
|  | Superiorparietal | Superiorparietal |
|  | Superiortemporal | Superiortemporal |
|  | Supramarginal | Supramarginal |
|  | Frontalpole | Frontalpole |
|  | Temporalpole | Temporalpole |
|  | Transversetemporal | Transversetemporal |
|  | Insula | Insula |

*Note: \* not bilateral*

### List of discovered loci

Please see Table S2 for an overview of the number of whole-genome significant SNPs, the number of independent SNPs, and the number of independent loci (see the Online Methods for definitions), per test and per feature set. The full tables of discovered loci for each feature set, for both MOSTest and min-P are in Data S1-S8. This includes information on lead SNP, genomic location, significance, and mapped genes, as outputted by FUMA.

**Table S2.** Number of whole-genome significant SNPs, independent SNPs and independent loci discovered, per test and per feature set.

| Test | Feature set | # significant SNPs | # independent SNPs | # loci discovered |
| --- | --- | --- | --- | --- |
| MOSTest | All | 93461 | 1425 | 347 |
| MOSTest | Subcortical | 46937 | 607 | 177 |
| MOSTest | Surface area | 33756 | 538 | 139 |
| MOSTest | Cortical thickness | 20443 | 229 | 71 |
| min-P | All | 24203 | 357 | 115 |
| min-P | Subcortical | 13194 | 228 | 90 |
| min-P | Surface area | 12650 | 182 | 53 |
| min-P | Cortical thickness | 9305 | 81 | 24 |

### Results from the replication

Table S3 lists the results from the replication attempt of the main GWAS via MOSTest and min-P. For this, we used a new batch of neuroimaging data of 4,884 healthy White European UKB participants that was released in October 2019, after we ran our primary analyses and made the results of this available via bioRxiv. This data was processed identically to the main sample and then analysed through MOSTest and min-P. We subsequently calculated the percentage of loci discovered in the main analyses, per test and per feature set, that was nominally significant in this additional sample. As can be seen in Table S3, for each combination of test and feature set, the replication rate was approximately 40%, indicating no major difference in the replication rates. From this it follows that the absolute number of loci replicating is threefold higher for MOSTest than for min-P.

**Table S3.**
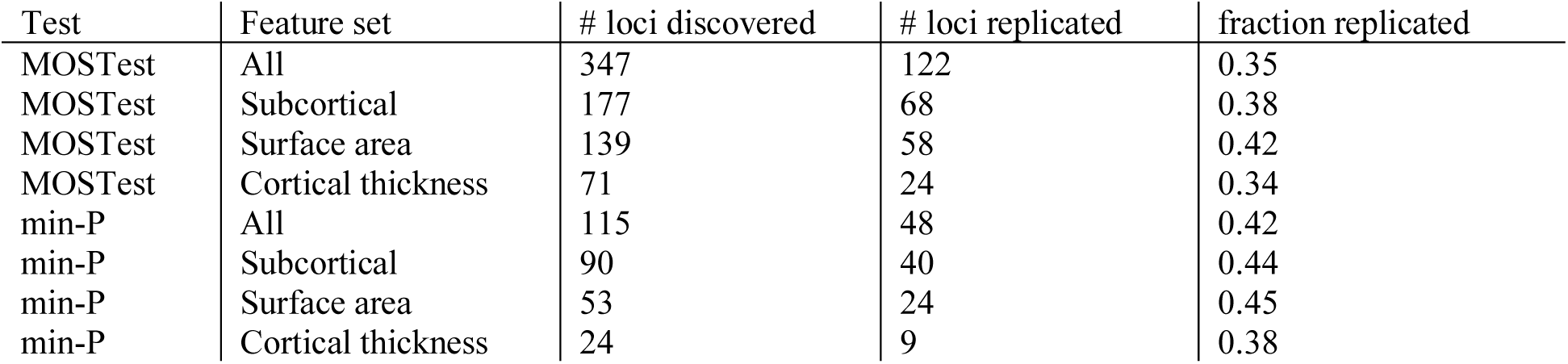
Results from the replication of the genome-wide significant loci, identified through MOSTest and min-P in an additional sample of 4,884 individuals. The column ‘# loci replicated’ indicates the amount of discovered loci in the main analysis that surpass a nominal significance (p=.05) in the replication sample. The ‘fraction replicated’ column divides the number of loci discovered in the main analysis with the number of loci that replicated.

| Test | Feature set | # loci discovered | # loci replicated | fraction replicated |
| --- | --- | --- | --- | --- |
| MOSTest | All | 347 | 122 | 0.35 |
| MOSTest | Subcortical | 177 | 68 | 0.38 |
| MOSTest | Surface area | 139 | 58 | 0.42 |
| MOSTest | Cortical thickness | 71 | 24 | 0.34 |
| min-P | All | 115 | 48 | 0.42 |
| min-P | Subcortical | 90 | 40 | 0.44 |
| min-P | Surface area | 53 | 24 | 0.45 |
| min-P | Cortical thickness | 24 | 9 | 0.38 |

### Validation and formal comparison between MOSTest and other tools

Current multivariate approaches, such as canonical correlation analysis as implemented in MV-PLINK^5^ and ordinal regression as implemented in MultiPhen^4^, perform a multiple regression in an opposite direction: the genotype vector is used as an outcome variable, while each phenotype is turned into an explanatory variable. The p-value is then calculated from an F-test, which tests for an association between the genotype vector and the most predictive linear combination of phenotypes at each SNP. Figure S1 compares the -log10(p-value), calculated by MV-PLINK and MOSTest. MV-PLINK takes 10,000x longer to run (requiring approximately 250K CPU hours, instead of 24 CPU hours with MOSTest), it was therefore infeasible to run the analysis on the entire set of 7.4M SNPs. Instead we tested a set of 356 LD-independent SNPs (a subset of all genome-wide significant SNPs remaining after LD-based clumping with r2=0.6 threshold) with a p-value from min-P below the genome-wide significance threshold. The results show very high correlation (r=0.9976) between MV-PLINK and MOSTest -log10(p-values), with median of 14.16 (MOSTest) versus 14.40 (MV-PLINK). Another recently developed multivariate test, aMAT^27^, uses the same test statistic as MOSTest, but after applying regularization (spectral filtering) to the correlation matrix R. In our data regularization wasn’t necessary, as the conditioning number was reasonably low, see Table S4, leading to a well-defined matrix R^-1^.

Another recently developed multivariate test, aMAT^10^, uses the same test statistic as MOSTest, but after applying regularization (spectral filtering) to the correlation matrix R. In our data regularization wasn’t necessary, as the conditioning number was reasonably low, see Extended Data Table 2, leading to a well-defined matrix R^-1^.

**Extended Data Figure 1.**
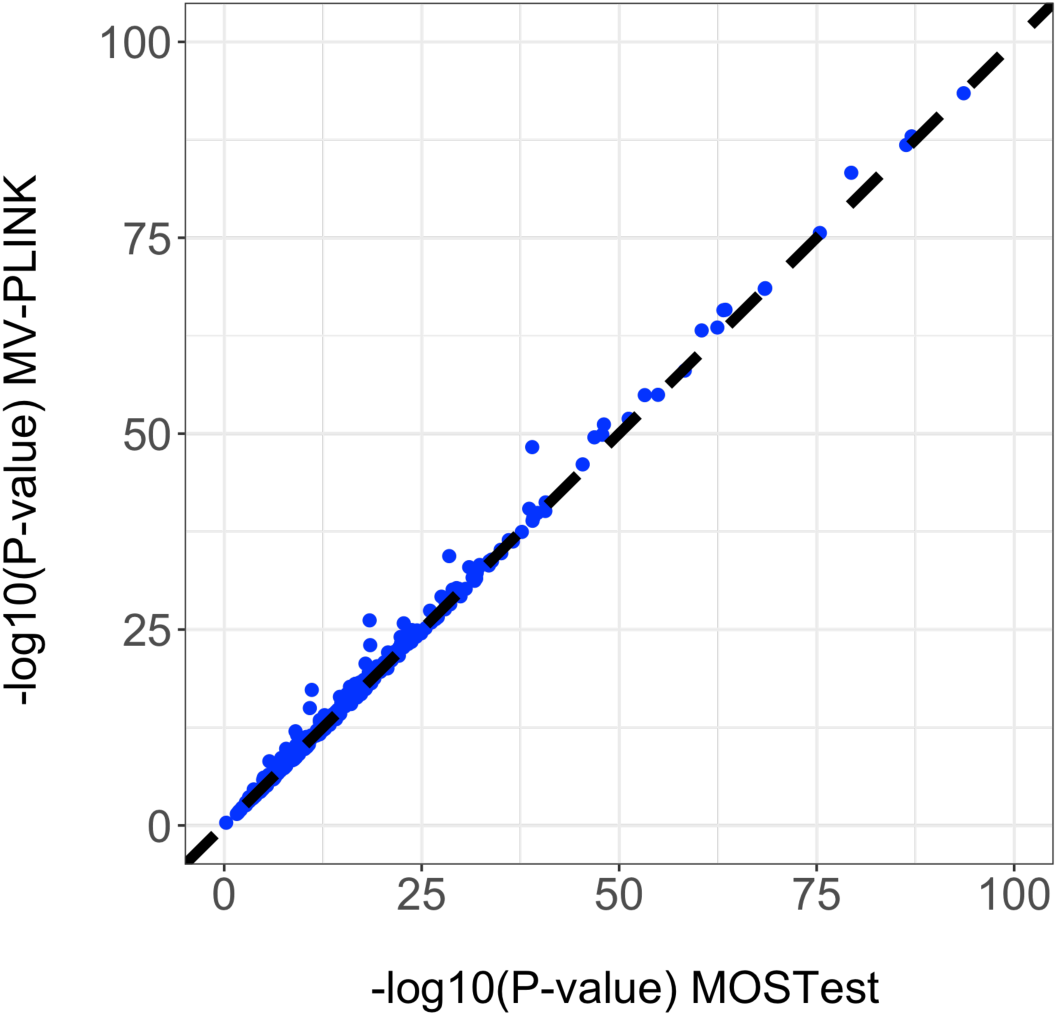
Comparison between MV-PLINK and MOSTest for a selected set of 356 SNPs, showing high correlation (r=0.9976) between MV-PLINK and MOSTest -log10(p-values).

**Table S4.** The conditioning number of the variance-covariance matrix (R), parameters of the Beta distribution (in min-P test) and Gamma distribution (in MOSTest).

| | #features | cond($R$) | Beta_a | Beta_b | Gamma_a | Gamma_b |
| --- | --- | --- | --- | --- | --- | --- |
| All | 171 | 2237.48 | 0.936 | 134.872 | 85.755 | 1.994 |
| Area | 68 | 1025.877 | 0.953 | 57.583 | 33.935 | 2.005 |
| Thickness | 68 | 191.599 | 0.938 | 53.641 | 34.049 | 1.998 |
| Subcortical | 35 | 132.63 | 0.913 | 20.977 | 17.484 | 2.002 |

### Rank-based inverse normal transformation

We carried out a rank-based inverse normal transformation of the measures, otherwise non-normally distributed measures can inflate p-values and thus elevate type-I error. Extended Data Figure 2 shows the empirical distribution of MOSTest and min-P test statistics under the null (calculated via permutations), along with p-value calculated from the test, with the rank-based INT, showing correct behaviour. Extended Data Figure 3 shows the distributions after running MOSTest without the transformation, leading to deviations, highlighting that this transformation is important for maintaining correct type-I error.

**Extended Data Figure 2.**
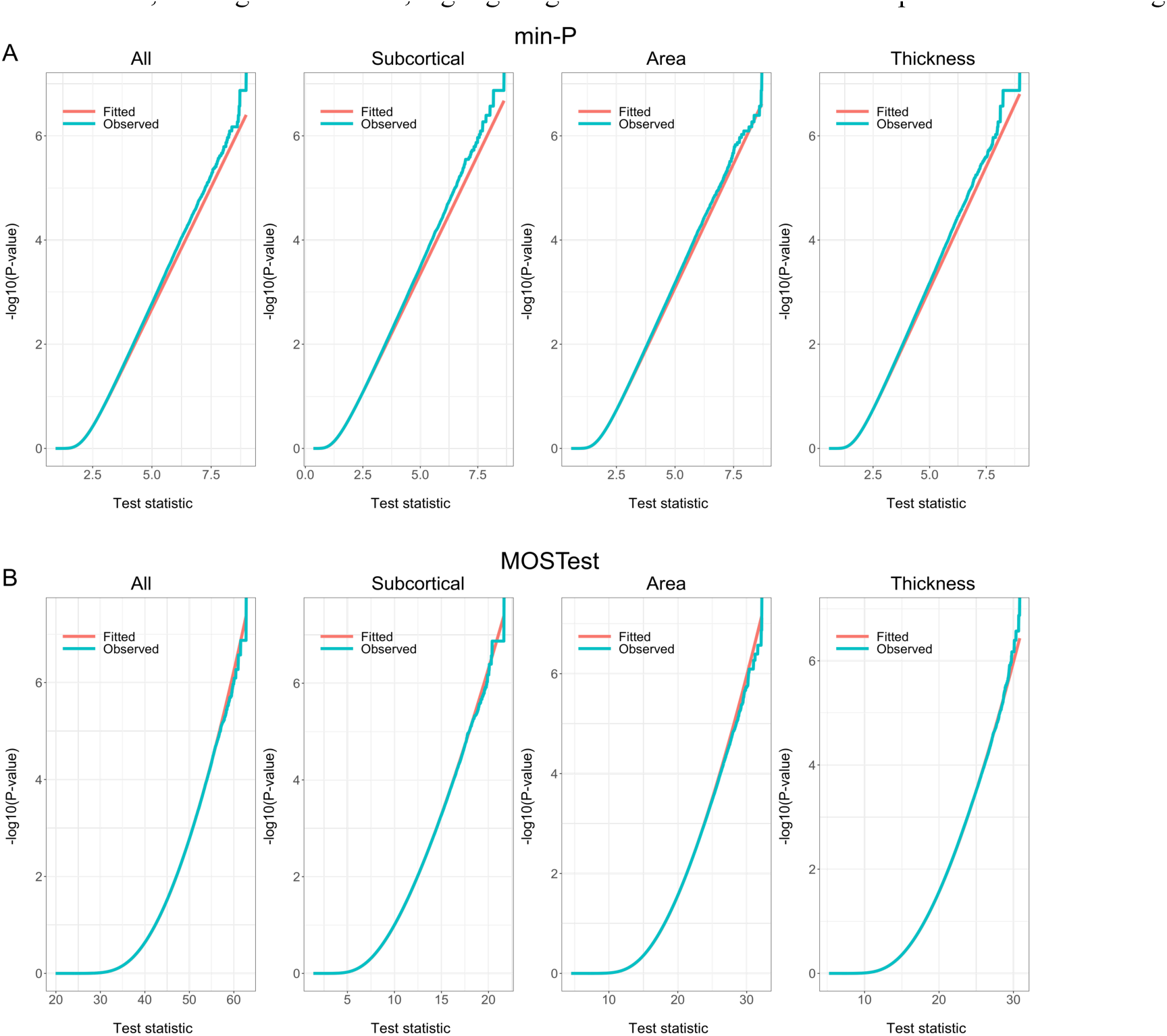
Each subplot shows an empirical distribution of the min-P (top row) and MOSTest (bottom row) test statistics under null (calculated via permutations), along with p-value calculated from the test. Columns correspond to different sets of phenotypes included in the analysis. OX axis show the actual value of the test statistic: - log10(min-P), for the min-P test, and z’ **R**^**-1**^ z for MOSTest. The “Observed” plot shows empirical distribution of the test statistic; “Fitted” plot shows p-values calculated from Gamma(a,b) distribution (MOSTest) and Beta(a,b) distribution (min-P) after fitting the two parameters to the observed data.

**Extended Data Figure 3.**
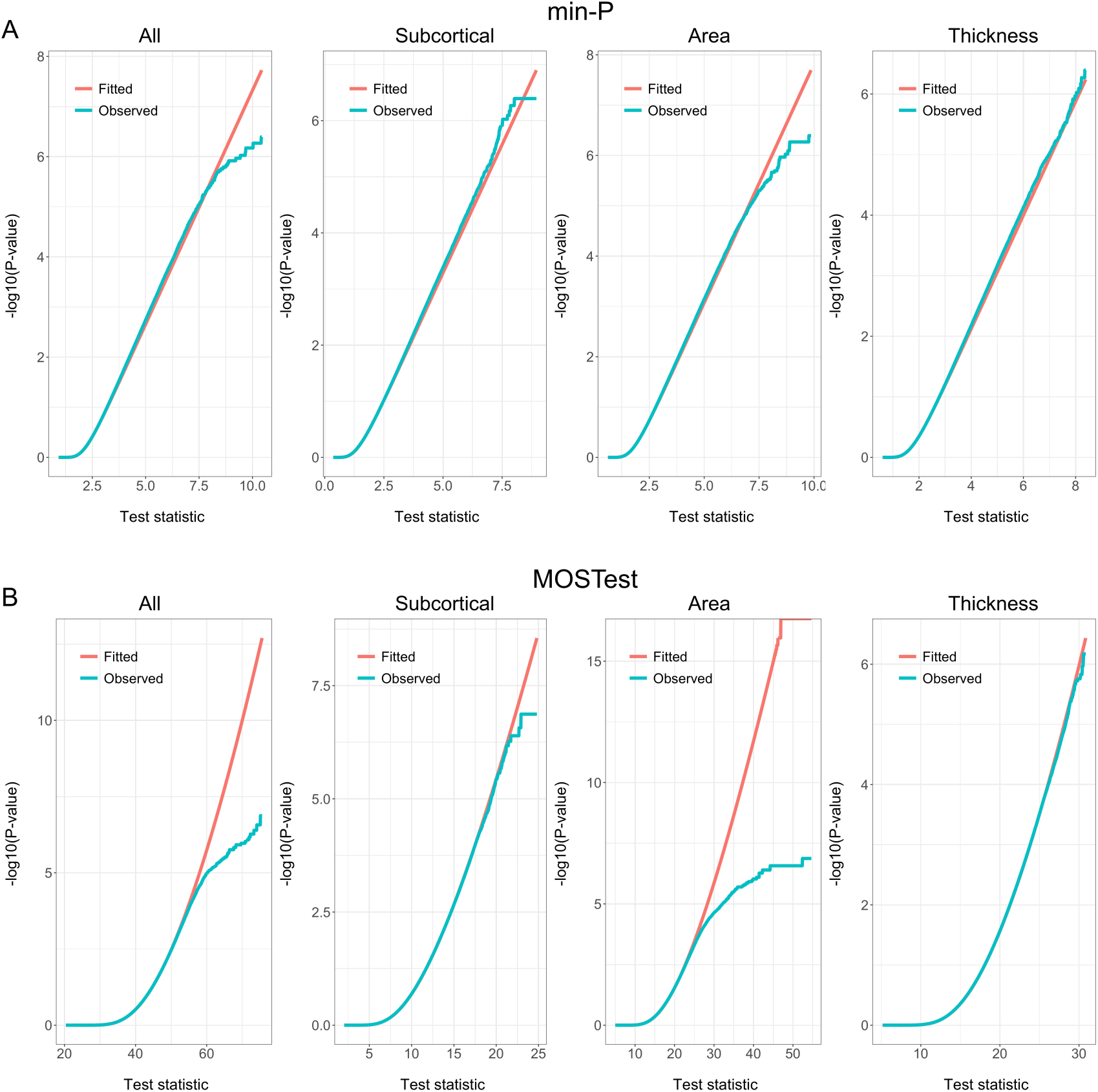
Distribution of min-P and MOSTest p-values under permutation, without applying rank-based inverse normal transformation, showing large deflection in the tails of the distribution. Appearance as in the previous figure.

### Simulations for validation of correct type-I error in the presence of polygenic signal

Extended Data Figure 2 shows that under permutation, i.e. in the absence of the genetic signal, MOSTest p-values are uniformly distributed. We performed further simulations on synthetic data to validate that MOSTest has correct type-I, not just under null, but also in the presence of a polygenic signal. We used the real genotype matrix of 26,502 individuals from our main analysis and simulated 175 phenotypes using a simple additive genetic model *y* = *Gβ* + *ϵ*, with the same phenotypic correlation as we observed in our combined set of feature (cortical area, cortical thickness and subcortical volumes, including CSF, left and right vessel, and optic chiasm). We randomly chose a set of 10K “causal” SNPs, but constrained them to odd chromosomes only, leaving even chromosomes free of genetic signal. 10K “causal” SNPs were shared across the 175 phenotypes, but each phenotype has its own randomly generated *β* vector of genetic effects. SNP heritability of each phenotype was h2=0.3. Applying MOSTest to test data, we observe the resulting signal was completely flat for even chromosomes, showing that regions that are not in LD with causal variants aren’t picked by MOSTest. These results are visualized in Extended Data Figure 4.

**Extended Data Figure 4.**
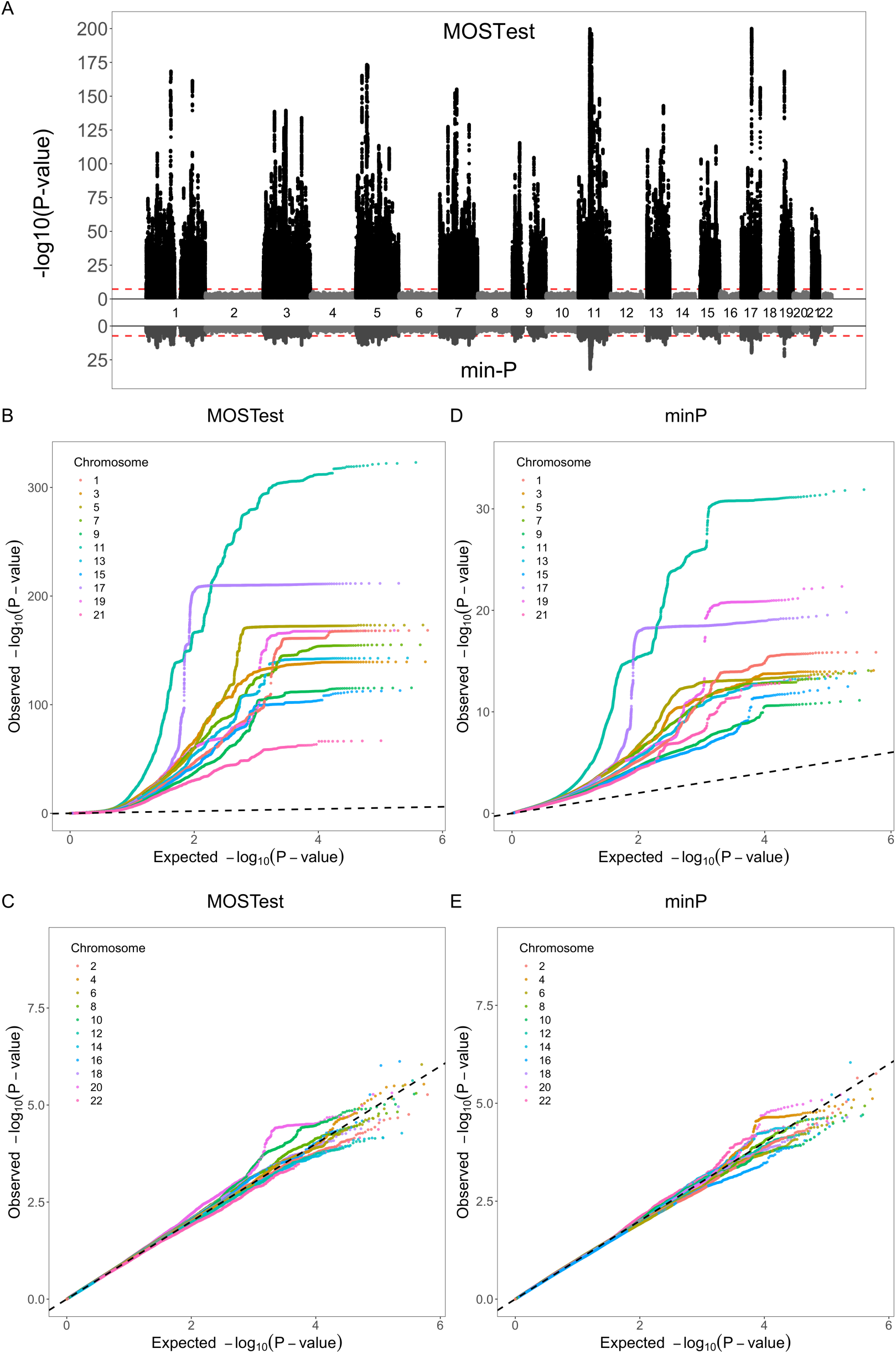
showing Manhattan and per-chromosome QQ plots for a synthetic dataset of 175 simulated phenotypes, with h=0.3 heritability, with 10K causal variants randomly assigned to odd chromosomes. No signal is observed for even chromosomes (as expected).

### Genomic inflation

We applied LD score regression^27^ to test for genomic inflation in MOSTest and min-P results. The results, listed in Extended Data Table 5, show no genomic inflation.

**Extended Data Table 5.** Genomic inflation analysis with LD Score Regression intercept (from partitioned LDSC 1kG phase3 reference), confirming no confounding effects (stratification, cryptic relatedness) in MOSTest p-values.

| Feature set | Test | LambdaGC | Intercept from LDSC |
| --- | --- | --- | --- |
| All | min-P | 1.6524 | 0.9864 (0.0096) |
| All | MOSTest | 2.0217 | 0.9247 (0.015) |
| Subcortical | min-P | 1.5511 | 1.0062 (0.0086) |
| Subcortical | MOSTest | 1.7179 | 0.9742 (0.0106) |
| Area | min-P | 1.453 | 0.9913 (0.0093) |
| Area | MOSTest | 1.5733 | 0.9748 (0.0112) |
| Thickness | min-P | 1.4034 | 1.0085 (0.0068) |
| Thickness | MOSTest | 1.5254 | 0.9952 (0.0085) |

**Extended Data Figure 5.**
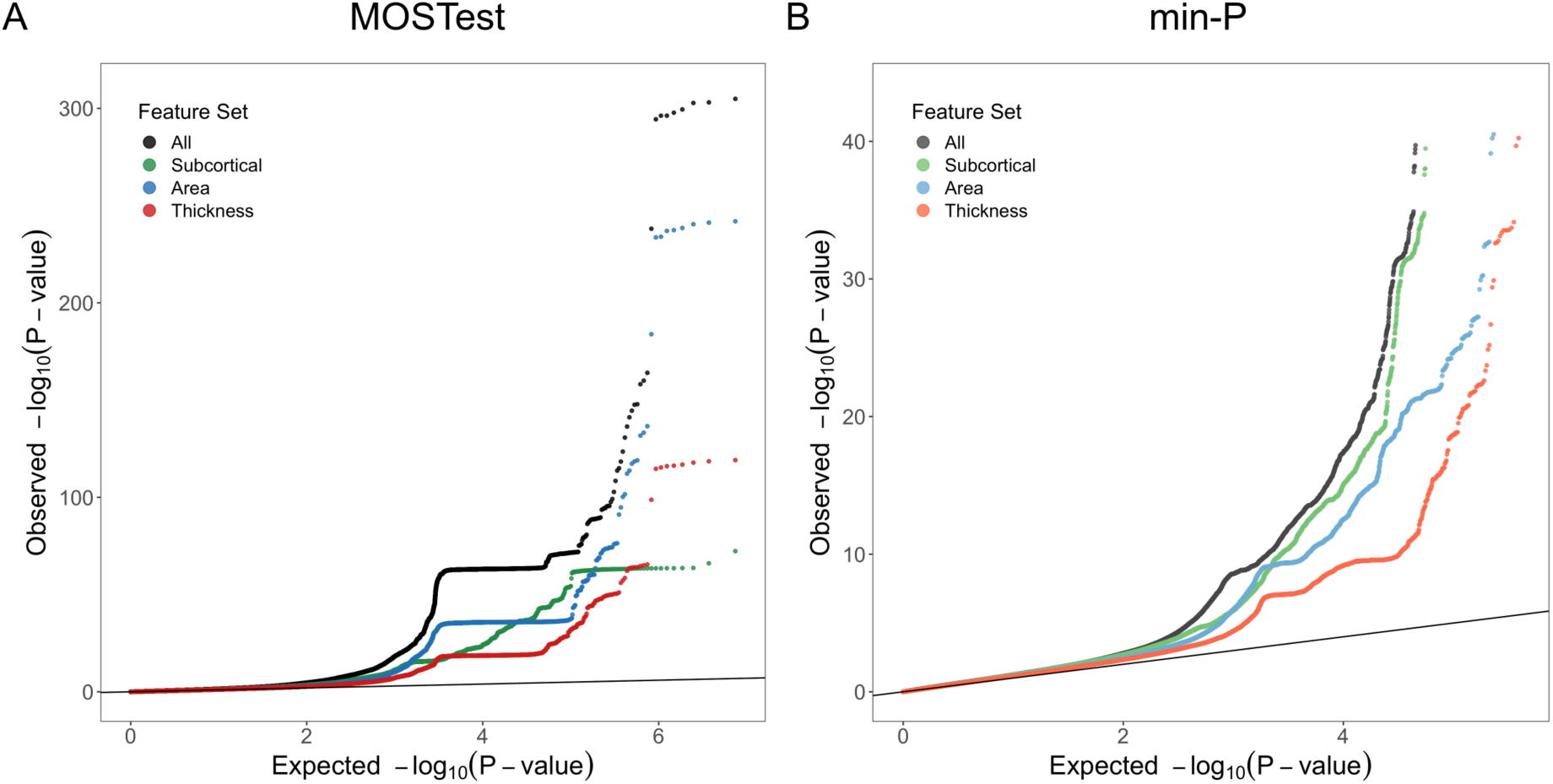
Quantile-Quantile plots for the p-values derived from MOSTest (A) and min-P (B).

## References

1. Grasby, K. L. et al. The genetic architecture of the human cerebral cortex. bioRxiv 399402 (2018). doi:10.1101/399402

2. Satizabal, C. L. et al. Genetic Architecture of Subcortical Brain Structures in Over 40,000 Individuals Worldwide. bioRxiv 173831 (2017).

3. Porter, H. F. & O’Reilly, P. F. Multivariate simulation framework reveals performance of multi-trait GWAS methods. Sci. Rep. 7, 38837 (2017).

4. O’Reilly, P. F. et al. MultiPhen: joint model of multiple phenotypes can increase discovery in GWAS. PLoS One 7, e34861 (2012).

5. Ferreira, M. A. R. & Purcell, S. M. A multivariate test of association. Bioinformatics 25, 132–133 (2008).

6. Alexander-Bloch, A., Giedd, J. N. & Bullmore, E. Imaging structural co-variance between human brain regions. Nat. Rev. Neurosci. 14, 322 (2013).

7. Panizzon, M. S. et al. Distinct genetic influences on cortical surface area and cortical thickness. Cereb. Cortex bhp026 (2009).

8. Frei, O. et al. Bivariate causal mixture model quantifies polygenic overlap between complex traits beyond genetic correlation. Nat. Commun. 10, 2417 (2019).

9. Van Der Sluis, S., Verhage, M., Posthuma, D. & Dolan, C. V. Phenotypic complexity, measurement bias, and poor phenotypic resolution contribute to the missing heritability problem in genetic association studies. PLoS One 5, e13929 (2010).

10. Wu, C. Multi-trait genome-wide analyses of the brain imaging phenotypes in UK Biobank. bioRxiv 758326 (2019). doi:10.1101/758326

11. Van der Sluis, S., Posthuma, D. & Dolan, C. V. TATES: efficient multivariate genotype-phenotype analysis for genome-wide association studies. PLoS Genet. 9, e1003235 (2013).

12. Fischl, B. et al. Whole brain segmentation: automated labeling of neuroanatomical structures in the human brain. Neuron 33, 341–355 (2002).

13. Desikan, R. S. et al. An automated labeling system for subdividing the human cerebral cortex on MRI scans into gyral based regions of interest. Neuroimage 31, 968–980 (2006).

14. Elliott, L. T. et al. Genome-wide association studies of brain imaging phenotypes in UK Biobank. Nature 562, 210 (2018).

15. Holland, D. et al. Beyond SNP Heritability: Polygenicity and Discoverability of Phenotypes Estimated with a Univariate Gaussian Mixture Model. bioRxiv 498550 (2019). doi:10.1101/498550

16. Chen, C.-H. et al. Hierarchical genetic organization of human cortical surface area. Science 335, 1634–1636 (2012).

17. Chen, C.-H. et al. Genetic topography of brain morphology. Proc. Natl. Acad. Sci. 110, 17089 LP – 17094 (2013).

18. Kong, X.-Z. et al. Mapping cortical brain asymmetry in 17,141 healthy individuals worldwide via the ENIGMA Consortium. Proc. Natl. Acad. Sci. 115, E5154–E5163 (2018).

19. de Leeuw, C. A., Mooij, J. M., Heskes, T. & Posthuma, D. MAGMA: generalized gene-set analysis of GWAS data. PLoS Comput. Biol. 11, e1004219 (2015).

20. Watanabe, K., Taskesen, E., Bochoven, A. & Posthuma, D. Functional mapping and annotation of genetic associations with FUMA. Nat. Commun. 8, 1826 (2017).

21. Andreassen, O. A. et al. Improved detection of common variants associated with schizophrenia and bipolar disorder using pleiotropy-informed conditional false discovery rate. PLoS Genet. 9, e1003455 (2013).

22. Watanabe, K. et al. A global overview of pleiotropy and genetic architecture in complex traits. Nat. Genet. 1–10 (2019).

23. Miller, K. L. et al. Multimodal population brain imaging in the UK Biobank prospective epidemiological study. Nat. Neurosci. 19, 1523–1536 (2016).

24. Sudlow, C. et al. UK biobank: an open access resource for identifying the causes of a wide range of complex diseases of middle and old age. PLoS Med. 12, e1001779 (2015).

25. Bycroft, C. et al. The UK Biobank resource with deep phenotyping and genomic data. Nature 562, 203–209 (2018).

26. Bensimhoun, M. N-dimensional cumulative function, and other useful facts about gaussians and normal densities. Jerusalem, Isr. Tech. Rep 1–8 (2009).

27. Finucane, H. K. et al. Partitioning heritability by functional annotation using genome-wide association summary statistics. Nat. Genet. 47, 1228 (2015).

