## Supplementary figures and images for "Making the MOSTest of imaging genetics"

### BrainMap101_rs6089075.png

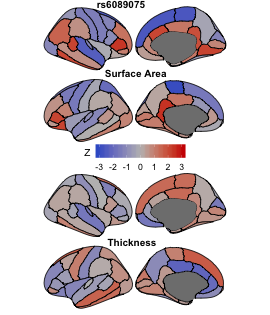

### BrainMap102_rs700059.png

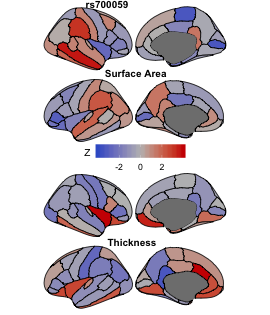

### BrainMap103_rs11607886.png

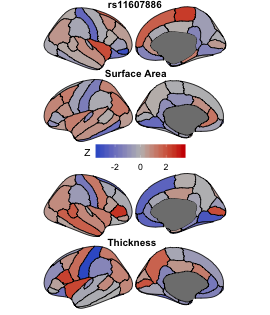

### BrainMap104_rs28456360.png

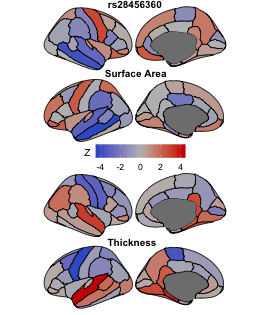

### BrainMap105_rs76244727.png

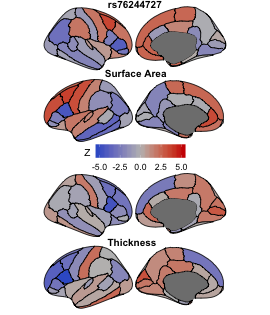

### BrainMap106_rs10797342.png

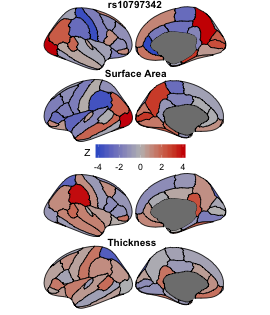

### BrainMap107_rs66716358.png

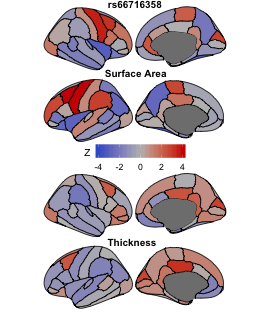

### BrainMap108_rs2332058.png

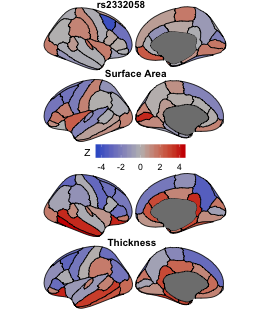

### BrainMap109_rs75925257.png

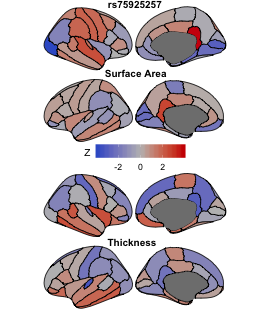

### BrainMap110_rs34254072.png

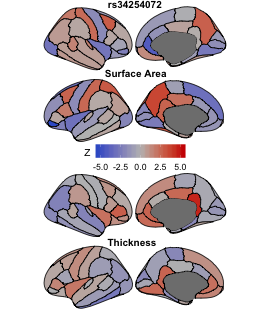

### BrainMap111_rs10283100.png

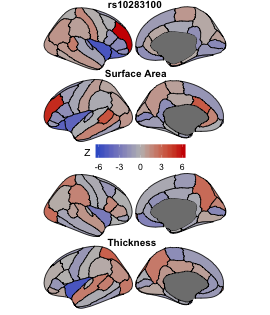

### BrainMap112_rs12460849.png

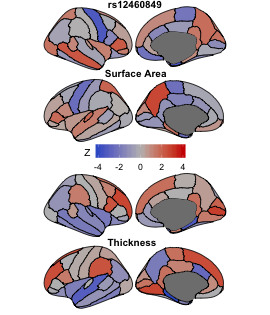

### BrainMap113_rs16948048.png

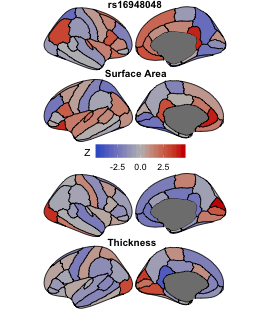

### BrainMap114_rs12025621.png

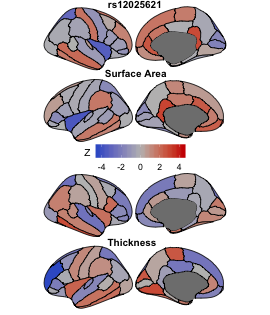

### BrainMap115_rs1058305.png

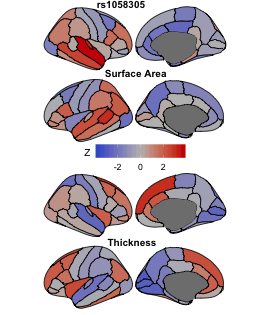

### BrainMap116_rs12708665.png

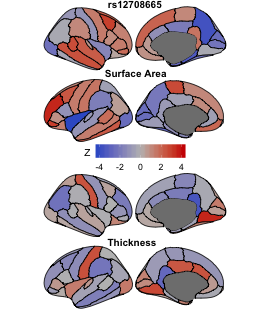

### BrainMap117_rs7782219.png

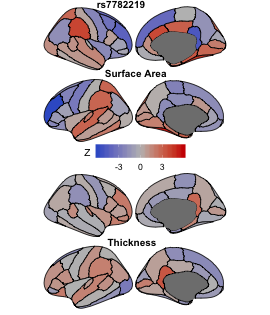

### BrainMap118_rs6799186.png

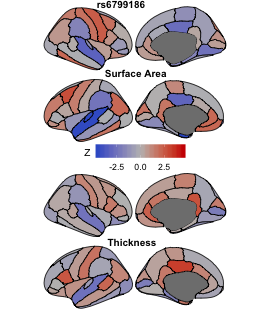

### BrainMap119_rs60900293.png

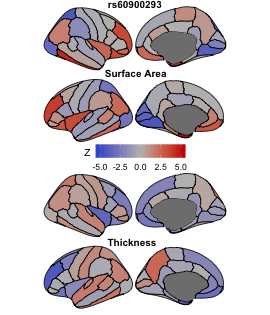

### BrainMap120_rs58419810.png

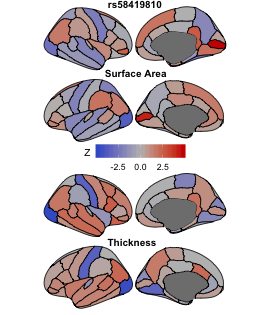

### BrainMap121_rs7095087.png

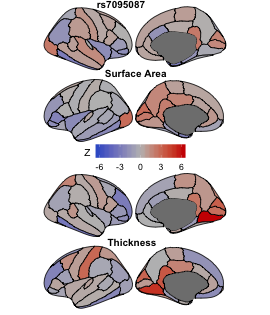

### BrainMap122_rs321406.png

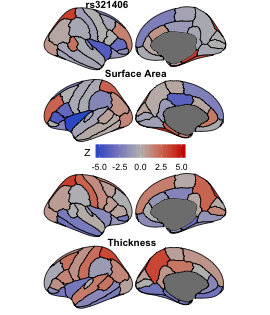

### BrainMap123_rs72766530.png

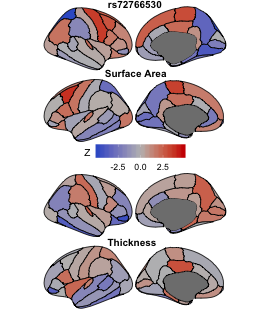

### BrainMap124_rs1654723.png

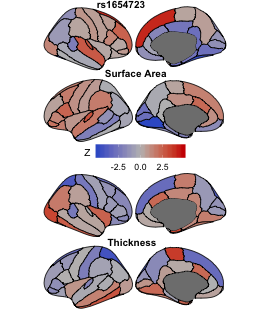

### BrainMap125_rs55732507.png

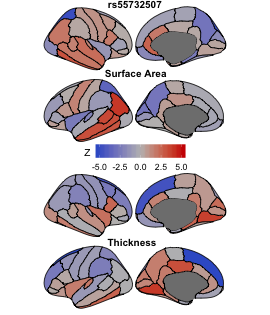

### BrainMap126_rs4507432.png

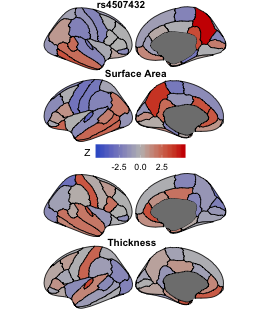

### BrainMap127_rs12767683.png

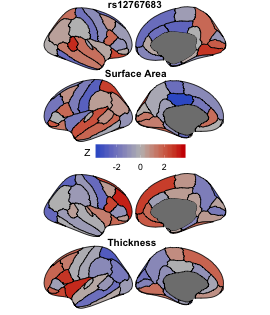

### BrainMap128_rs10861957.png

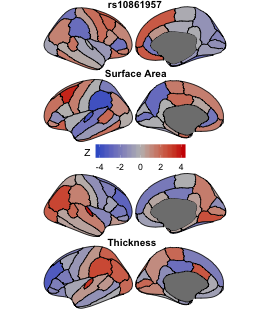

### BrainMap129_rs78428340.png

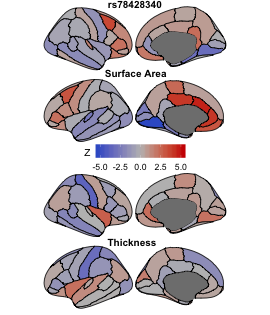

### BrainMap130_rs3012434.png

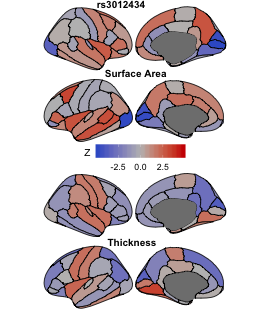

### BrainMap131_rs9870741.png

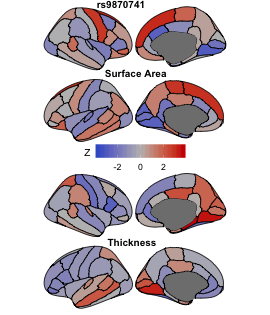

### BrainMap132_rs11243145.png

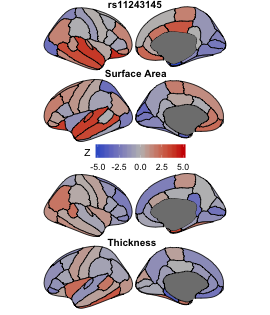

### BrainMap133_rs12028814.png

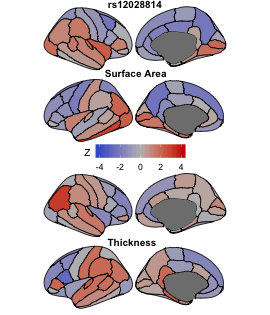

### BrainMap134_rs10037418.png

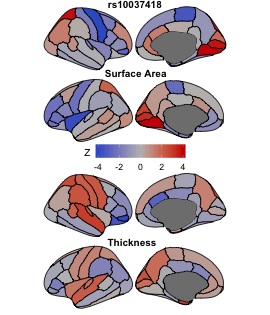

### BrainMap135_rs11181939.png

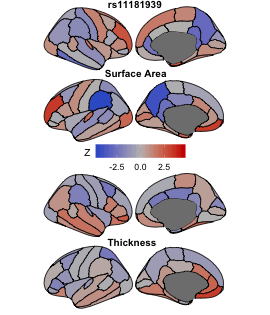

### BrainMap136_rs73058498.png

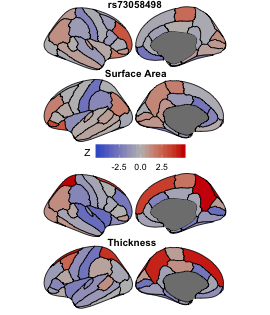

### BrainMap137_rs179744.png

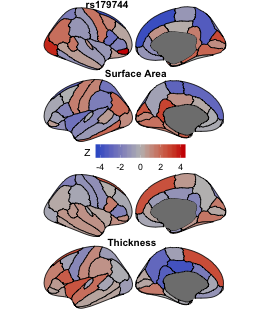

### BrainMap138_rs1413536.png

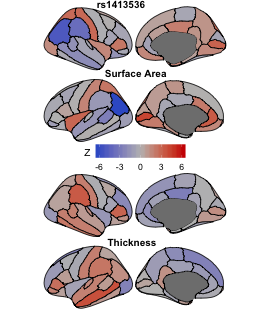

### BrainMap139_rs72794538.png

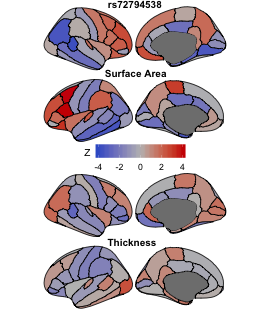

### BrainMap140_rs6757486.png

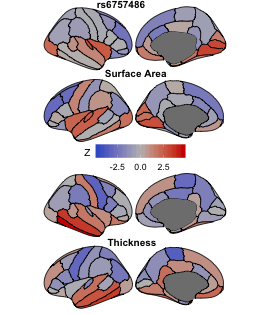

### BrainMap141_rs368928614.png

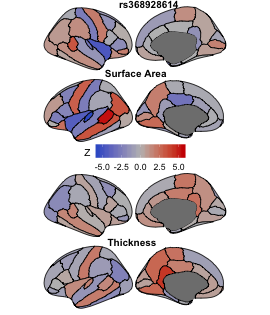

### BrainMap142_rs6557643.png

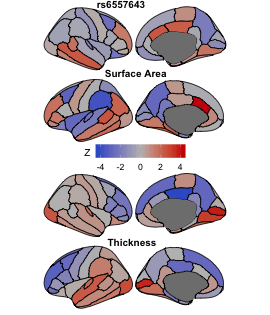

### BrainMap143_rs3743748.png

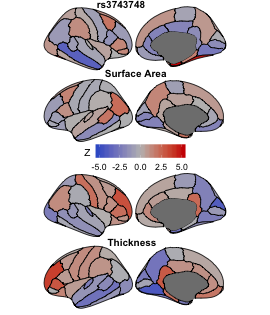

### BrainMap144_rs7601767.png

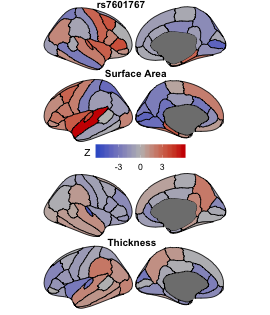

### BrainMap145_rs2664299.png

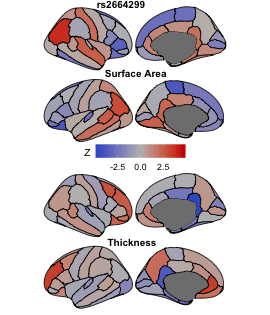

### BrainMap146_rs1069219.png

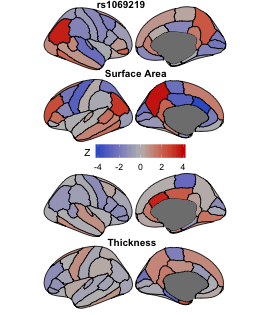

### BrainMap147_rs1073990.png

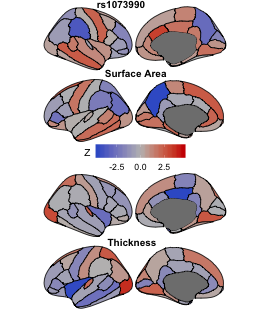

### BrainMap148_rs28472955.png

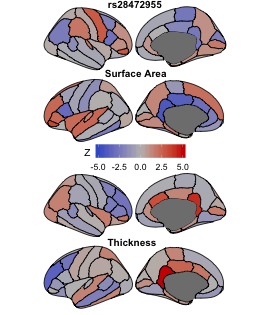

### BrainMap149_rs1536304.png

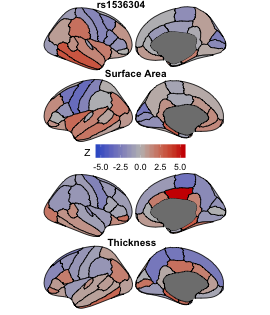

### BrainMap150_rs1694379.png

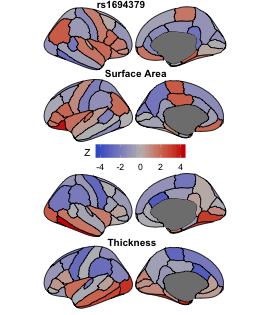

### BrainMap151_rs10239980.png

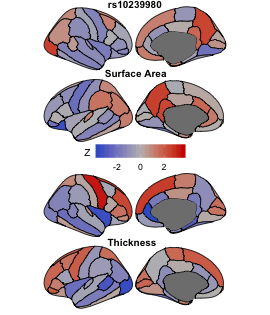

### BrainMap152_rs16841904.png

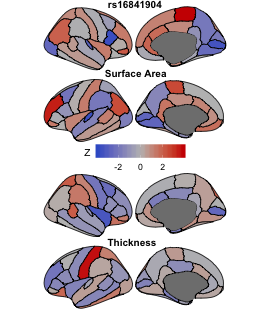

### BrainMap153_rs6723768.png

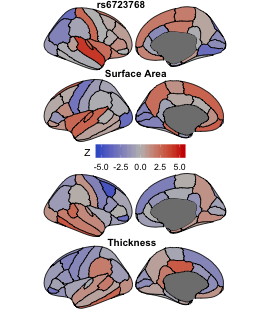

### BrainMap154_rs10149229.png

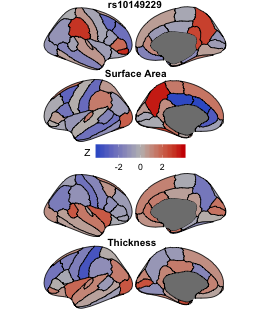

### BrainMap155_rs7808766.png

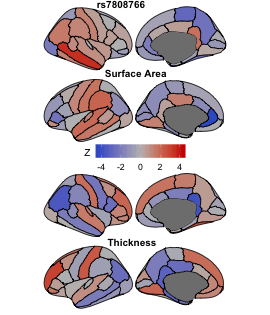

### BrainMap156_rs1481012.png

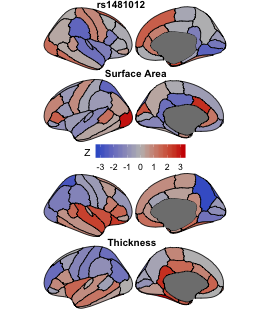

### BrainMap157_rs839996.png

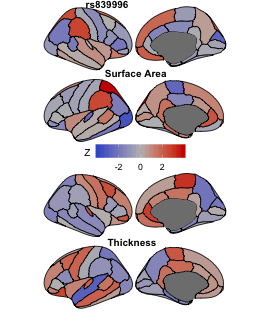

### BrainMap158_rs7214750.png

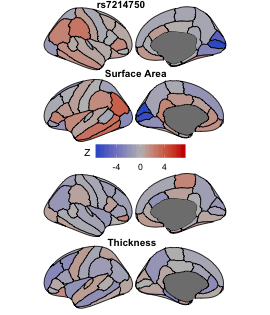

### BrainMap159_rs11727676.png

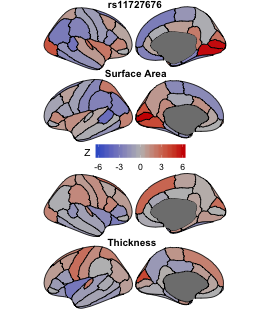

### BrainMap160_rs55985553.png

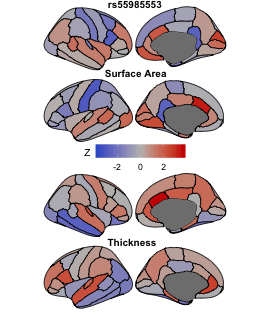

### BrainMap161_rs80323638.png

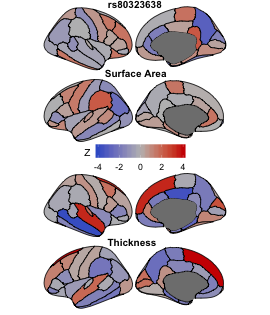

### BrainMap162_rs4128507.png

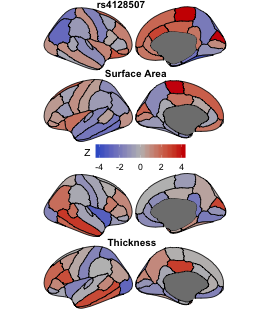

### BrainMap163_rs7586641.png

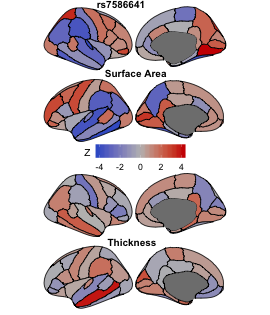

### BrainMap164_rs13246286.png

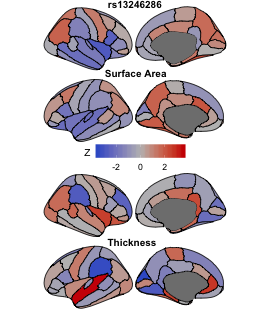

### BrainMap165_rs61741508.png

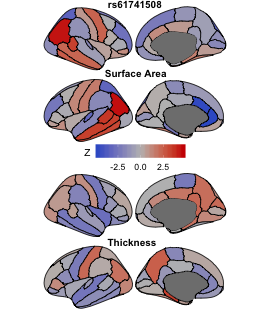

### BrainMap166_rs4530517.png

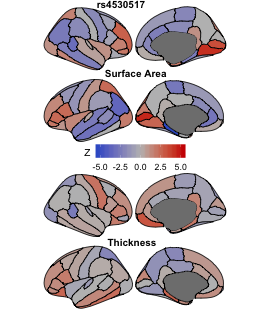

### BrainMap167_rs10835060.png

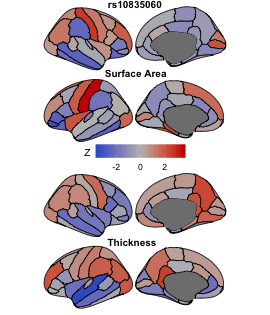

### BrainMap168_rs4683992.png

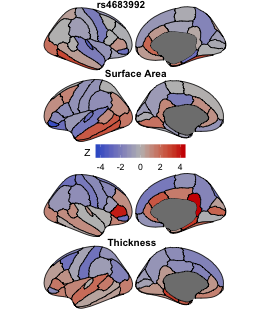

### BrainMap169_rs13079368.png

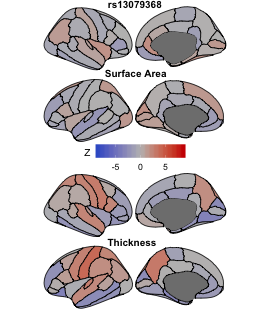

### BrainMap170_rs11683268.png

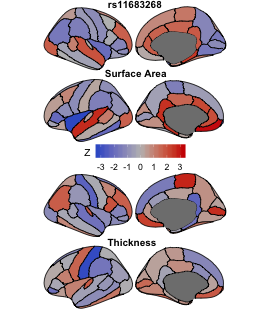

### BrainMap171_rs1476096.png

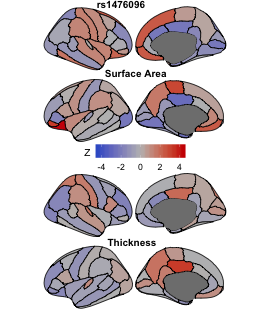

### BrainMap172_rs35772238.png

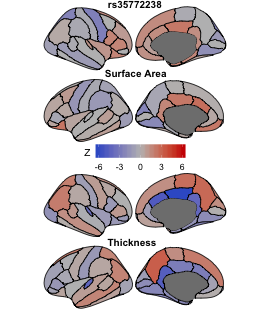

### BrainMap173_rs34722008.png

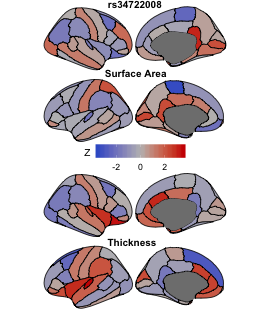

### BrainMap174_rs10229224.png

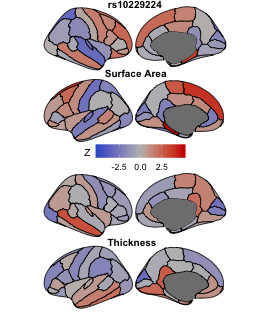

### BrainMap175_rs76880201.png

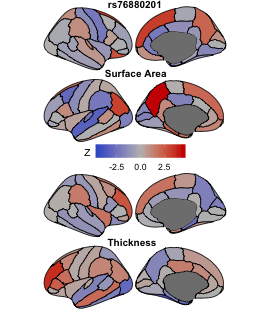

### BrainMap176_rs5027597.png

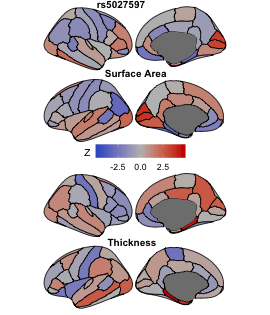

### BrainMap177_rs2167177.png

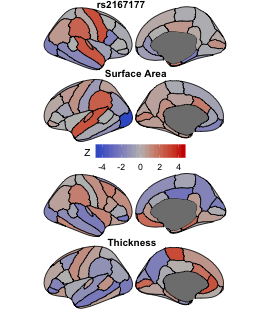

### BrainMap178_rs11206132.png

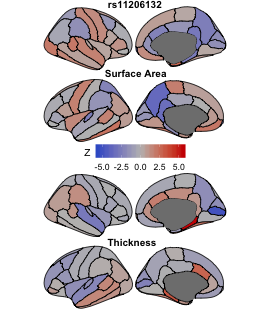

### BrainMap179_rs7956514.png

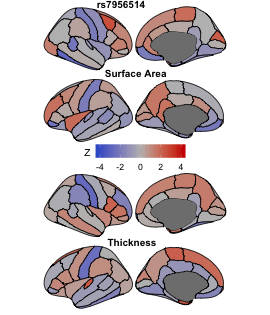

### BrainMap180_rs62365476.png

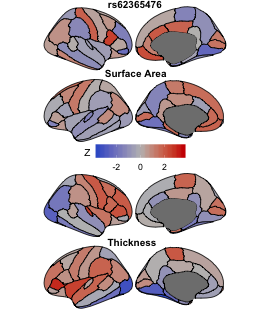

### BrainMap181_rs62381644.png

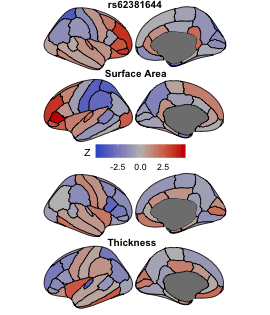

### BrainMap182_rs7968440.png

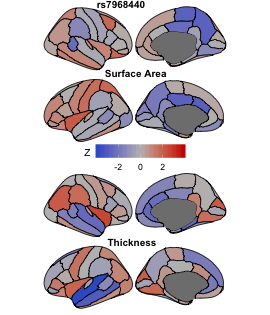

### BrainMap183_rs9809852.png

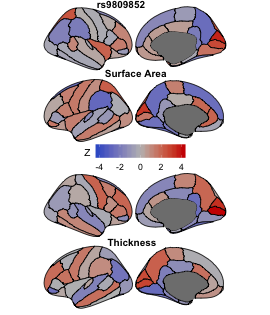

### BrainMap184_rs79272390.png

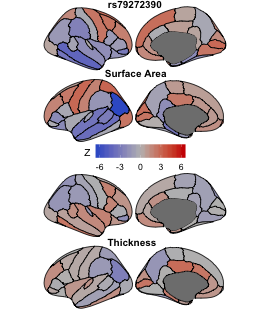

### BrainMap185_rs62509329.png

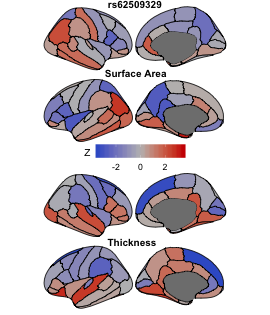

### BrainMap186_rs174538.png

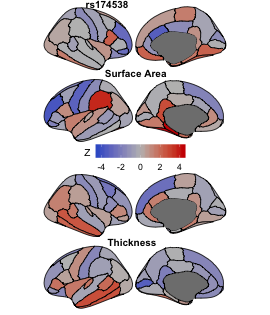

### BrainMap187_rs8179684.png

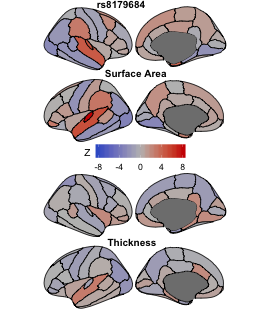

### BrainMap188_rs12373006.png

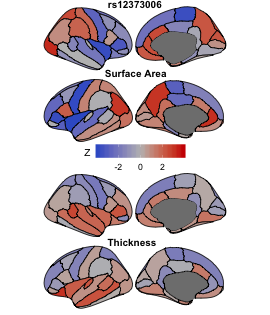

### BrainMap189_rs9926320.png

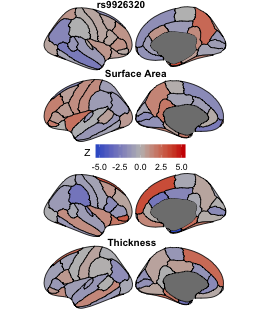

### BrainMap190_rs2415142.png

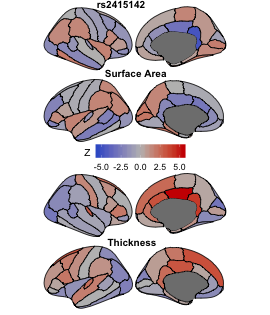

### BrainMap191_rs669952.png

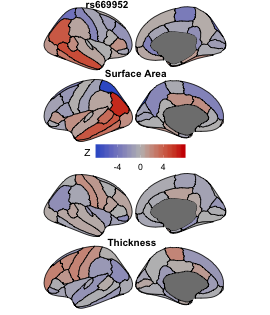

### BrainMap192_rs2474720.png

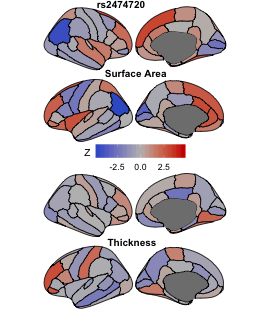

### BrainMap193_rs10821083.png

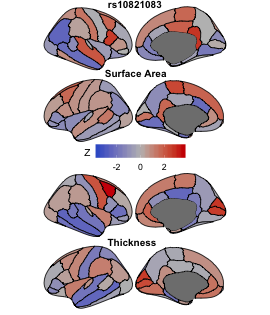

### BrainMap194_rs11135939.png

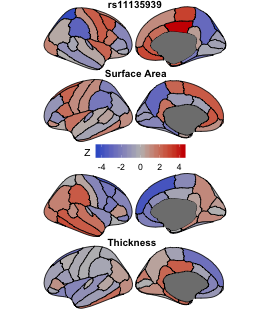

### BrainMap195_rs6426786.png

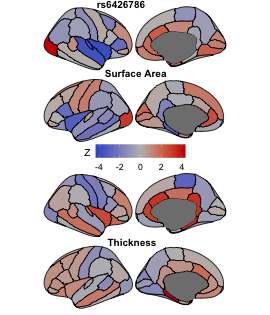

### BrainMap196_rs11022130.png

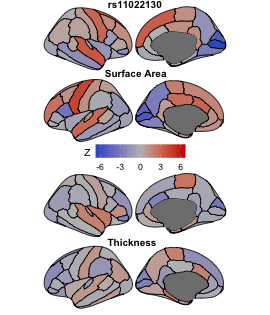

### BrainMap197_rs7017252.png

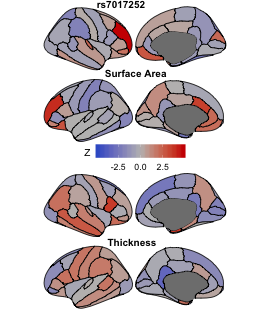

### BrainMap198_rs6087990.png

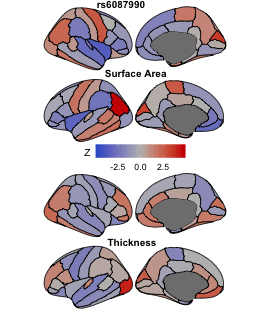

### BrainMap199_rs2296172.png

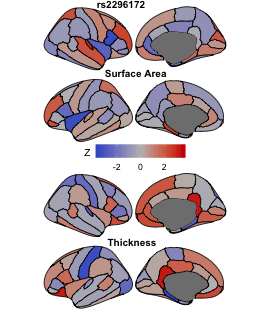

### BrainMap200_rs11955417.png

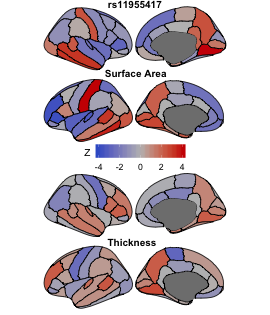
