## Supplementary figures and images for "Making the MOSTest of imaging genetics"

### BrainMap001_rs1080066.png

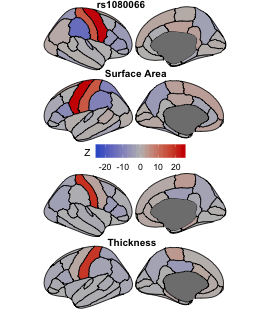

### BrainMap002_rs13107325.png

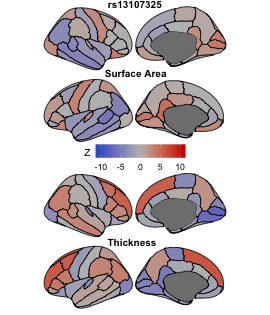

### BrainMap003_rs6658111.png

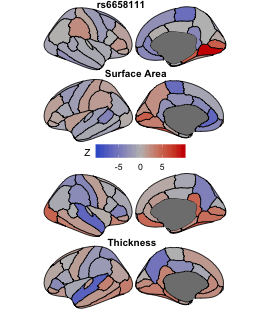

### BrainMap004_rs9937293.png

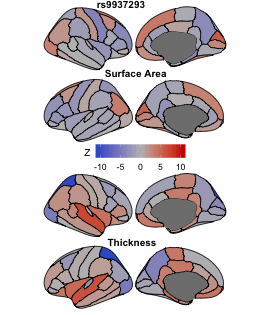

### BrainMap005_rs2279829.png

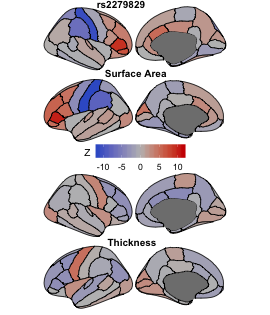

### BrainMap006_rs74826997.png

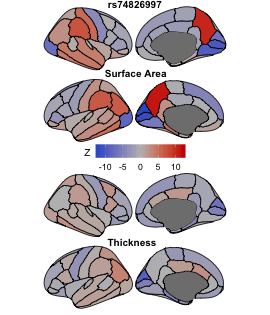

### BrainMap007_rs1909960.png

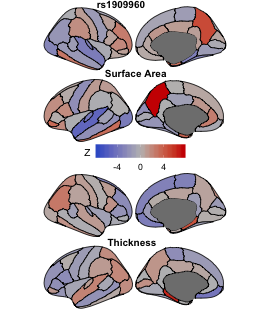

### BrainMap008_rs2532423.png

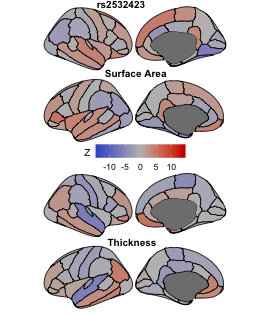

### BrainMap009_rs11245347.png

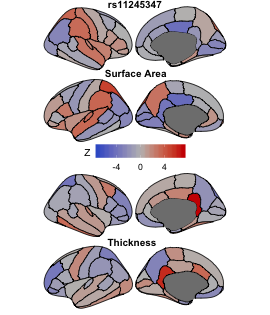

### BrainMap010_rs160459.png

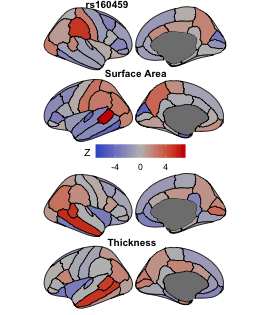

### BrainMap011_rs12146713.png

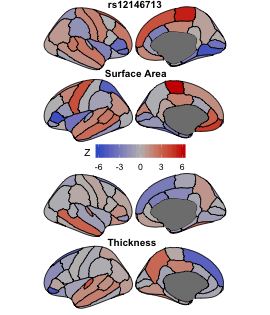

### BrainMap012_rs10878269.png

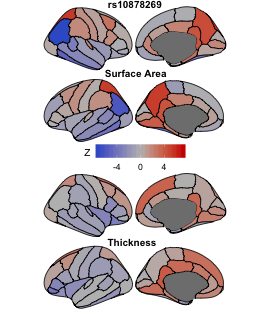

### BrainMap013_rs62132522.png

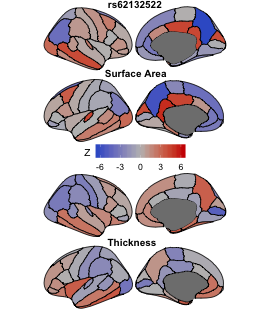

### BrainMap014_rs35124509.png

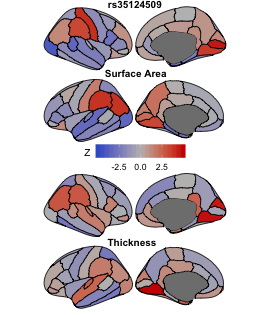

### BrainMap015_rs10939879.png

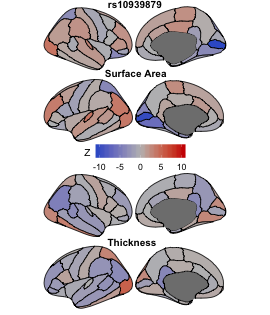

### BrainMap016_rs8053536.png

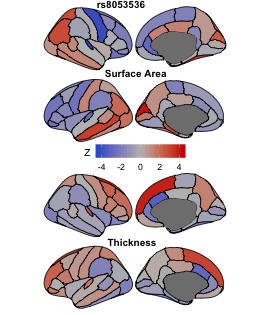

### BrainMap017_rs1490384.png

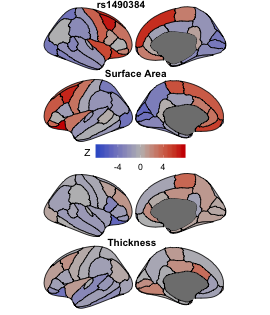

### BrainMap018_rs11111090.png

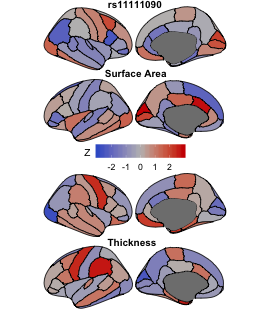

### BrainMap019_rs72754248.png

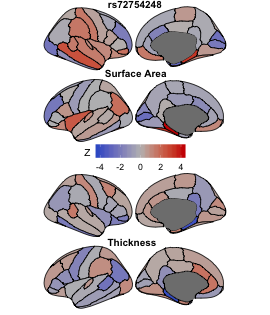

### BrainMap020_rs58237393.png

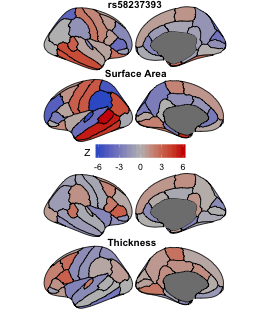

### BrainMap021_rs146607495.png

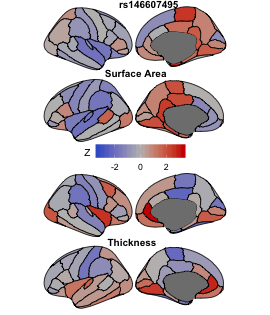

### BrainMap022_rs7303747.png

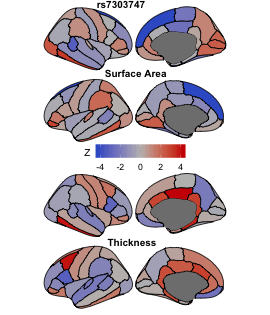

### BrainMap023_rs151057105.png

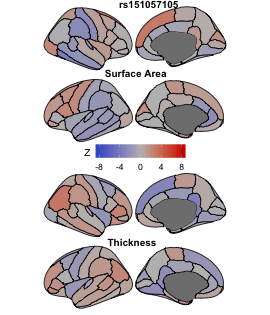

### BrainMap024_rs4721802.png

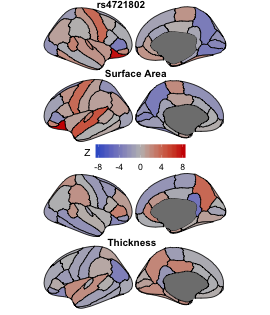

### BrainMap025_rs13723.png

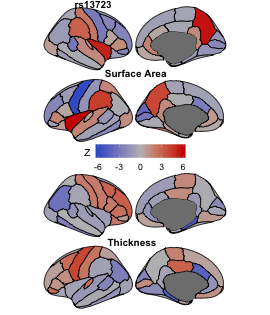

### BrainMap026_rs41298373.png

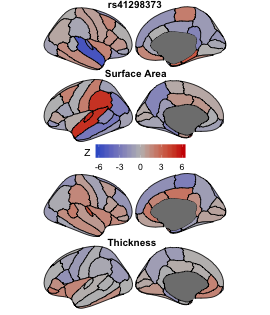

### BrainMap027_rs12256551.png

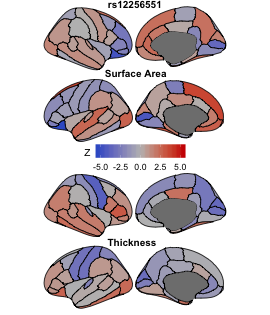

### BrainMap028_rs7764016.png

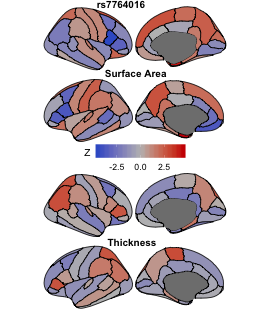

### BrainMap029_rs12700003.png

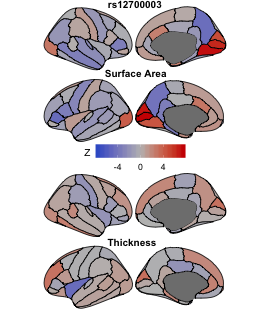

### BrainMap030_rs1639109.png

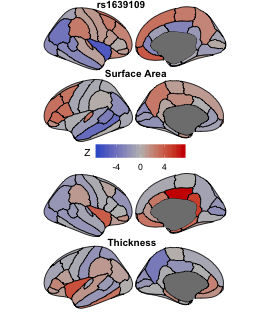

### BrainMap031_rs7005363.png

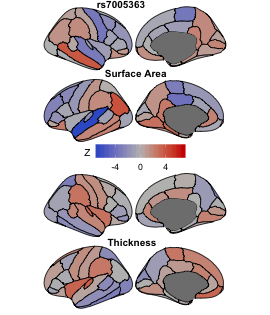

### BrainMap032_rs62348889.png

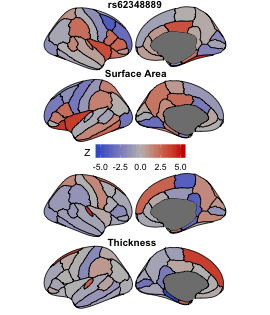

### BrainMap033_rs201650162.png

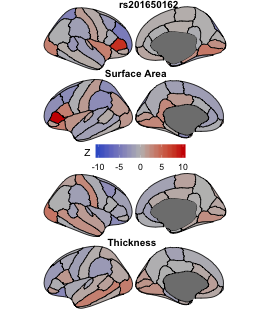

### BrainMap034_rs12769006.png

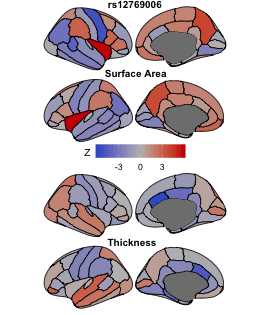

### BrainMap035_rs7023465.png

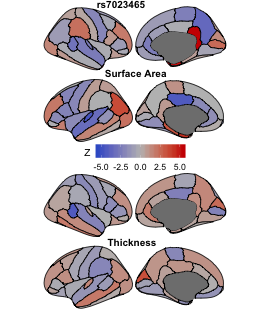

### BrainMap036_rs10512248.png

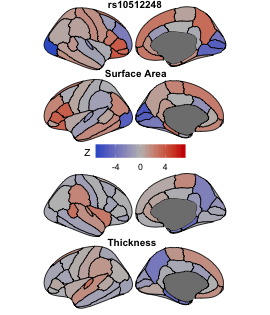

### BrainMap037_rs9345125.png

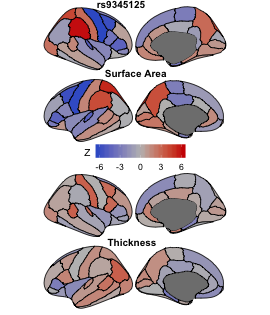

### BrainMap038_rs4842266.png

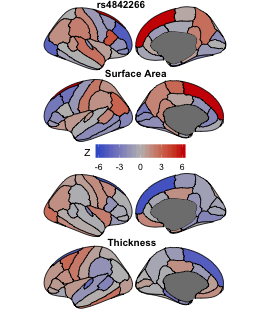

### BrainMap039_rs6510338.png

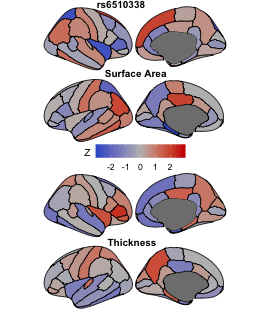

### BrainMap040_rs13430823.png

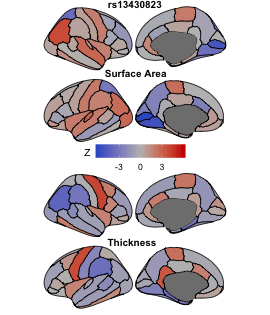

### BrainMap041_rs840585.png

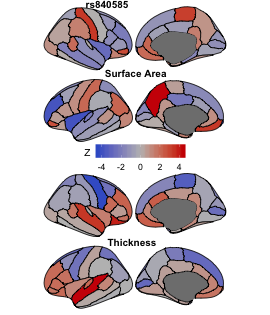

### BrainMap042_rs12526675.png

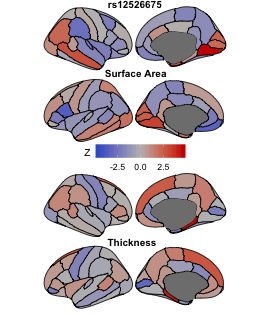

### BrainMap043_rs4663669.png

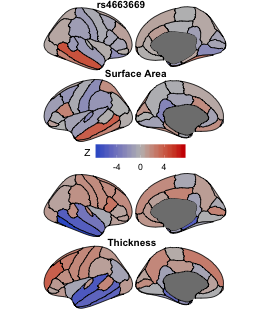

### BrainMap044_rs6126985.png

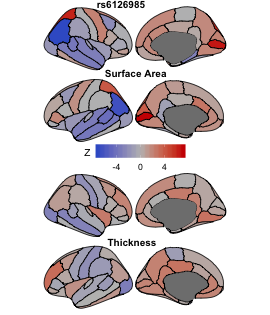

### BrainMap045_rs73006822.png

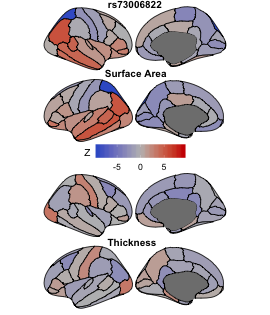

### BrainMap046_rs7562708.png

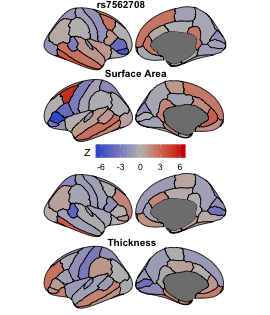

### BrainMap047_rs947934.png

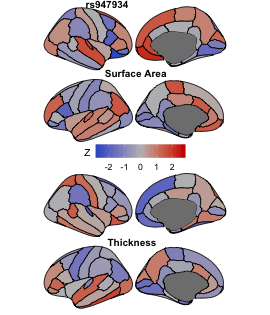

### BrainMap048_rs8287.png

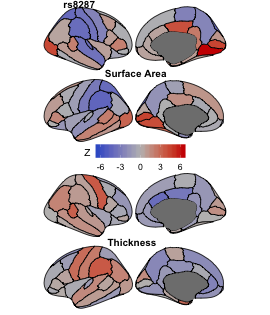

### BrainMap049_rs1285841.png

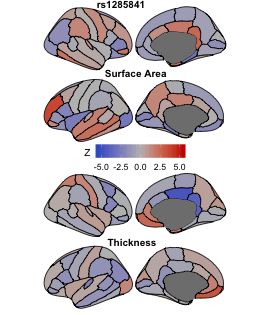

### BrainMap050_rs11206344.png

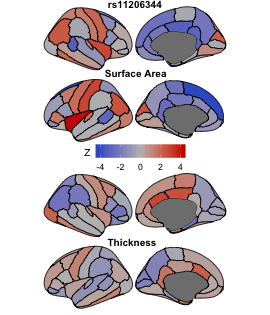

### BrainMap051_rs11669334.png

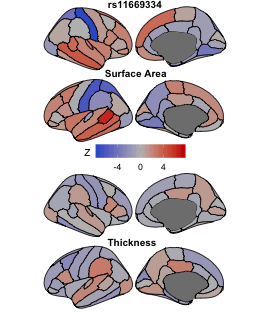

### BrainMap052_rs746698.png

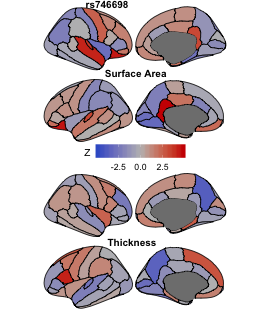

### BrainMap053_rs823080.png

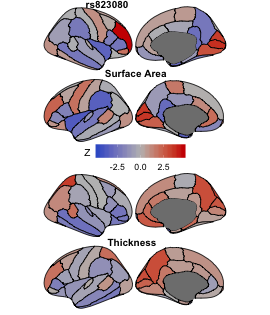

### BrainMap054_rs7531398.png

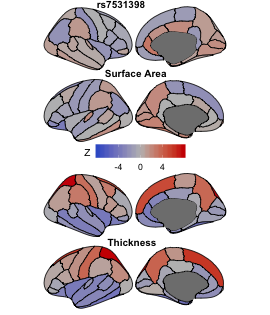

### BrainMap055_rs6806605.png

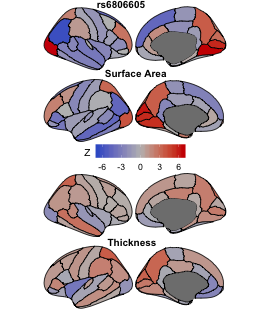

### BrainMap056_rs3850873.png

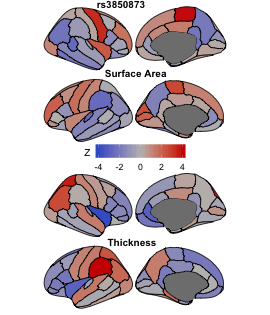

### BrainMap057_rs9545549.png

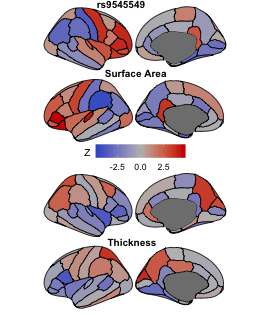

### BrainMap058_rs224331.png

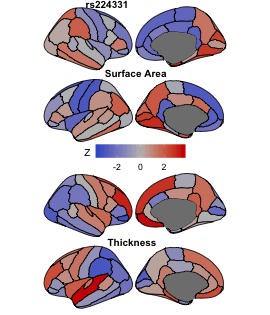

### BrainMap059_rs1451533.png

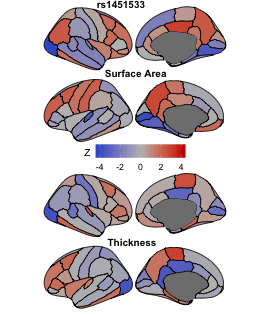

### BrainMap060_rs72786903.png

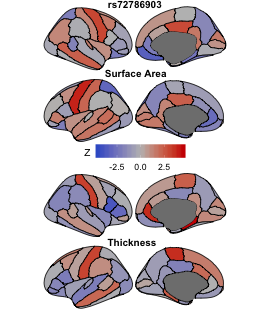

### BrainMap061_rs490340.png

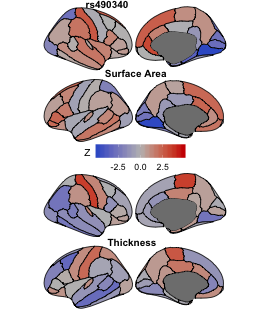

### BrainMap062_rs1693560.png

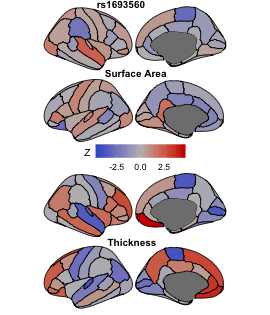

### BrainMap063_rs6133987.png

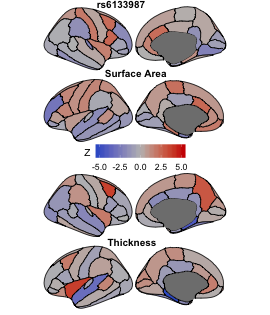

### BrainMap064_rs798497.png

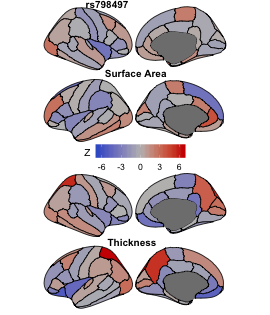

### BrainMap065_rs310528.png

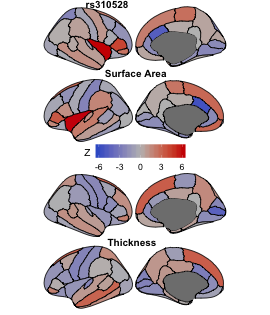

### BrainMap066_rs11674165.png

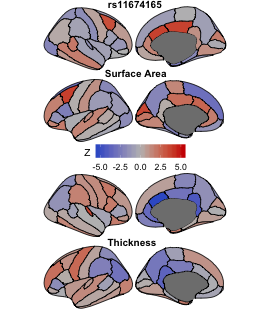

### BrainMap067_rs8012377.png

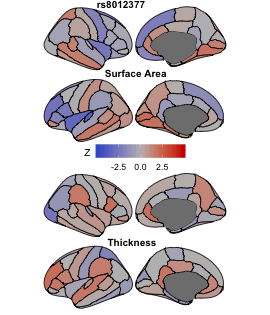

### BrainMap068_rs2678915.png

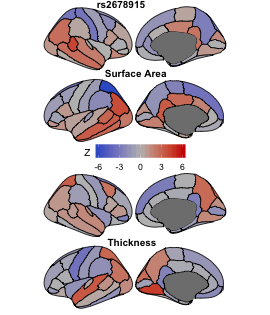

### BrainMap069_rs8042779.png

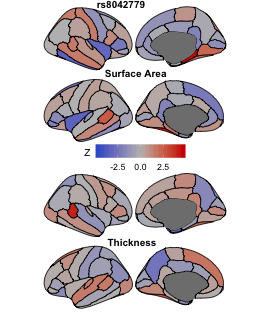

### BrainMap070_rs4703894.png

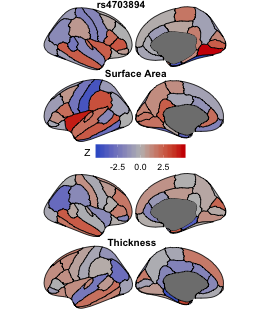

### BrainMap071_rs11126784.png

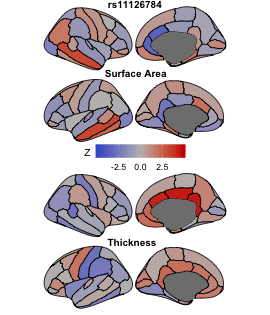

### BrainMap072_rs113057113.png

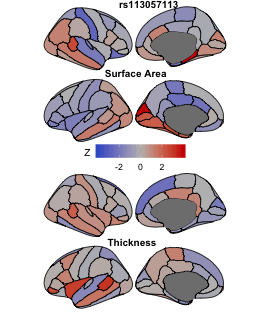

### BrainMap073_rs12997266.png

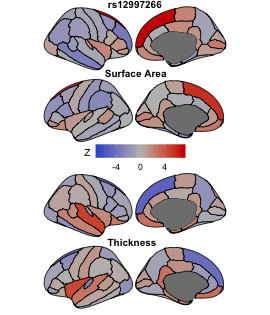

### BrainMap074_rs12322111.png

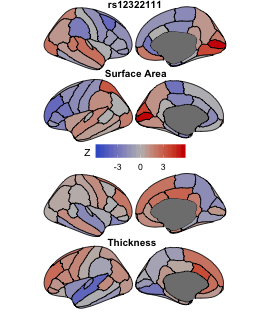

### BrainMap075_rs12408663.png

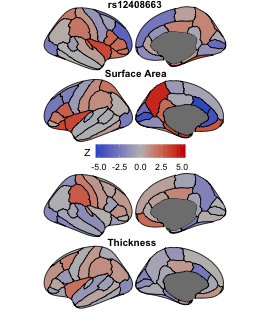

### BrainMap076_rs9954838.png

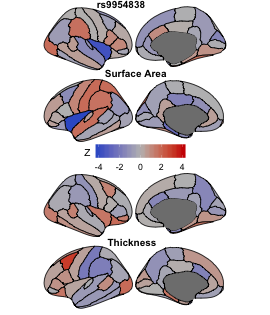

### BrainMap077_rs2790103.png

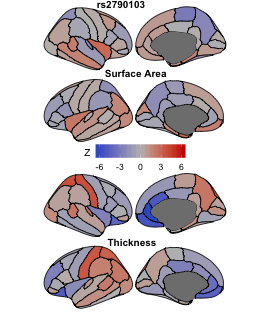

### BrainMap078_rs1520.png

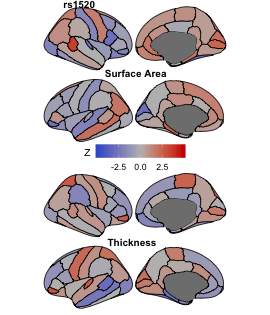

### BrainMap079_rs1915146.png

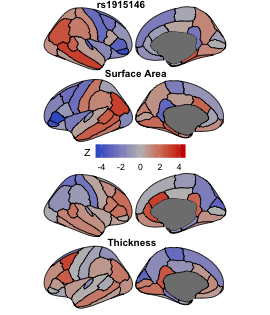

### BrainMap080_rs62086046.png

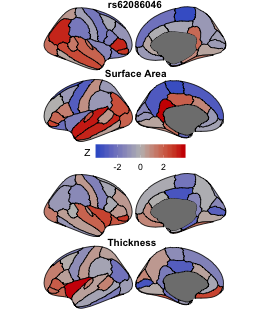

### BrainMap081_rs888812.png

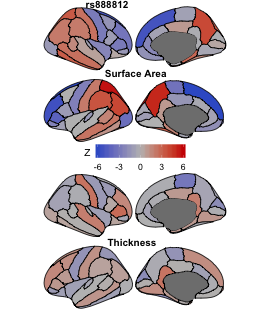

### BrainMap082_rs4391701.png

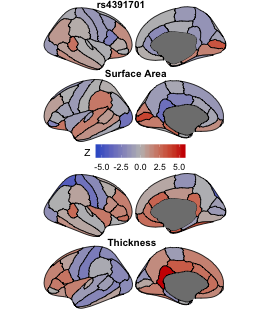

### BrainMap083_rs23595.png

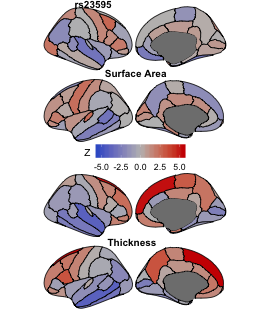

### BrainMap084_rs11652757.png

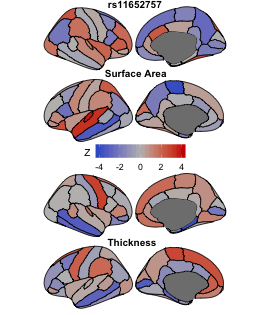

### BrainMap085_rs4128399.png

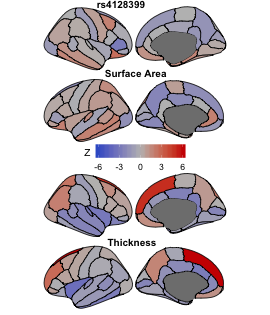

### BrainMap086_rs111398992.png

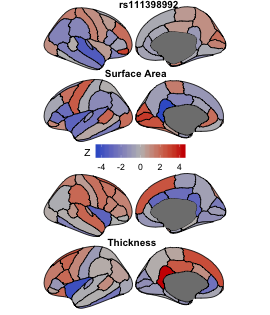

### BrainMap087_rs9534439.png

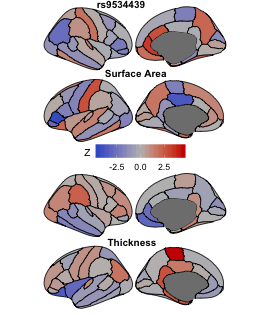

### BrainMap088_rs3847528.png

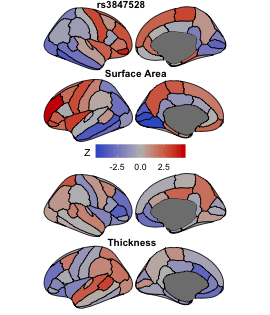

### BrainMap089_rs816329.png

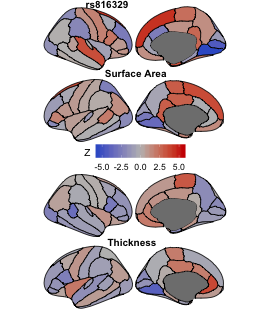

### BrainMap090_rs35614773.png

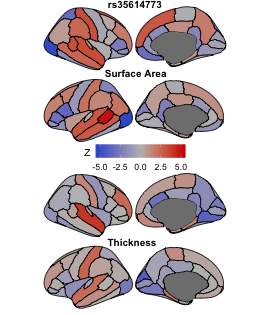

### BrainMap091_rs863216.png

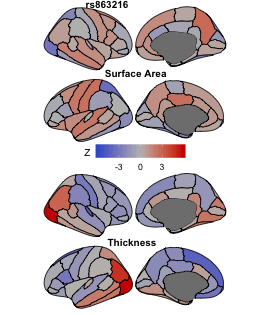

### BrainMap092_rs9376354.png

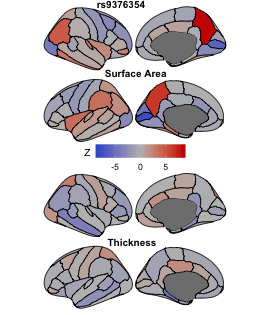

### BrainMap093_rs10237233.png

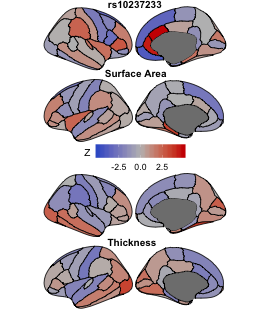

### BrainMap094_rs4398584.png

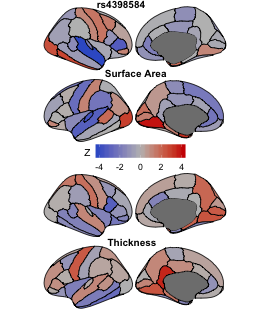

### BrainMap095_rs35061193.png

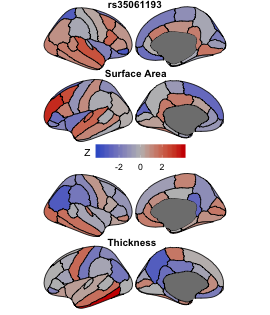

### BrainMap096_rs13150272.png

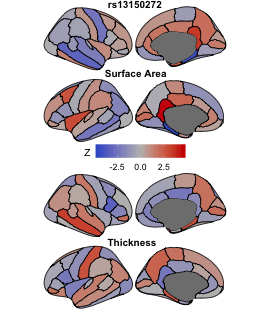

### BrainMap097_rs651858.png

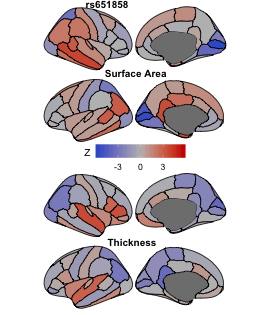

### BrainMap098_rs35447133.png

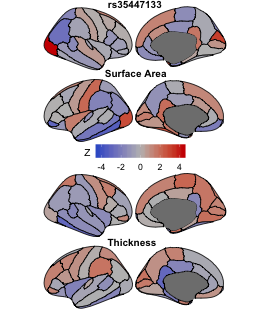

### BrainMap099_rs35891966.png

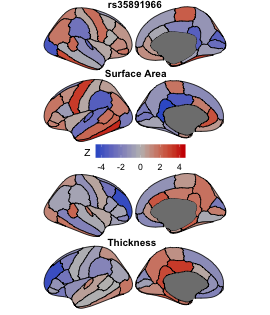

### BrainMap100_rs4811602.png

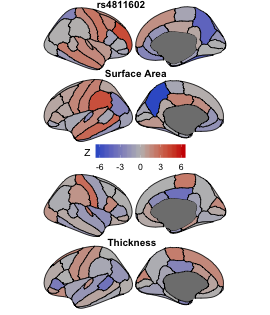
